# Visual-cortex-projecting neurons in mouse auditory cortex show distinct audiovisual responses independent of motor-related activity

**DOI:** 10.64898/2026.10.29.757229

**Authors:** Sietse Kiewiet, Umberto Olcese, Cyriel M.A. Pennartz

## Abstract

Cells in mouse primary visual cortex (V1) not only respond to visual stimulation, but also to sound. Part of this auditory modulation likely originates from auditory cortex (AC). Yet, the circuit-level functionality of the AC-to-V1 projection remains poorly understood. Here we performed acute in vivo high-density recordings in mouse AC during auditory, visual, and audiovisual stimulation, identifying AC-to-V1 projecting neurons by optogenetic tagging. [Compared to non-tagged neurons, AC-to-V1 neurons more often responded to auditory, visual, or audiovisual stimuli. Furthermore, most AC-to-V1 neurons responded to specific sound frequencies; a modest but significant fraction was selective for visual orientations. These results could not be explained by motor responses or single-neuron variations in firing rate, cortical recording depth, or waveform type. These findings show that AC-to-V1 neurons carry sensory-specific signals, as opposed to nonspecific modulations, and exhibit distinctive response profiles compared to non-tagged AC neurons.

**Graphical abstract:** 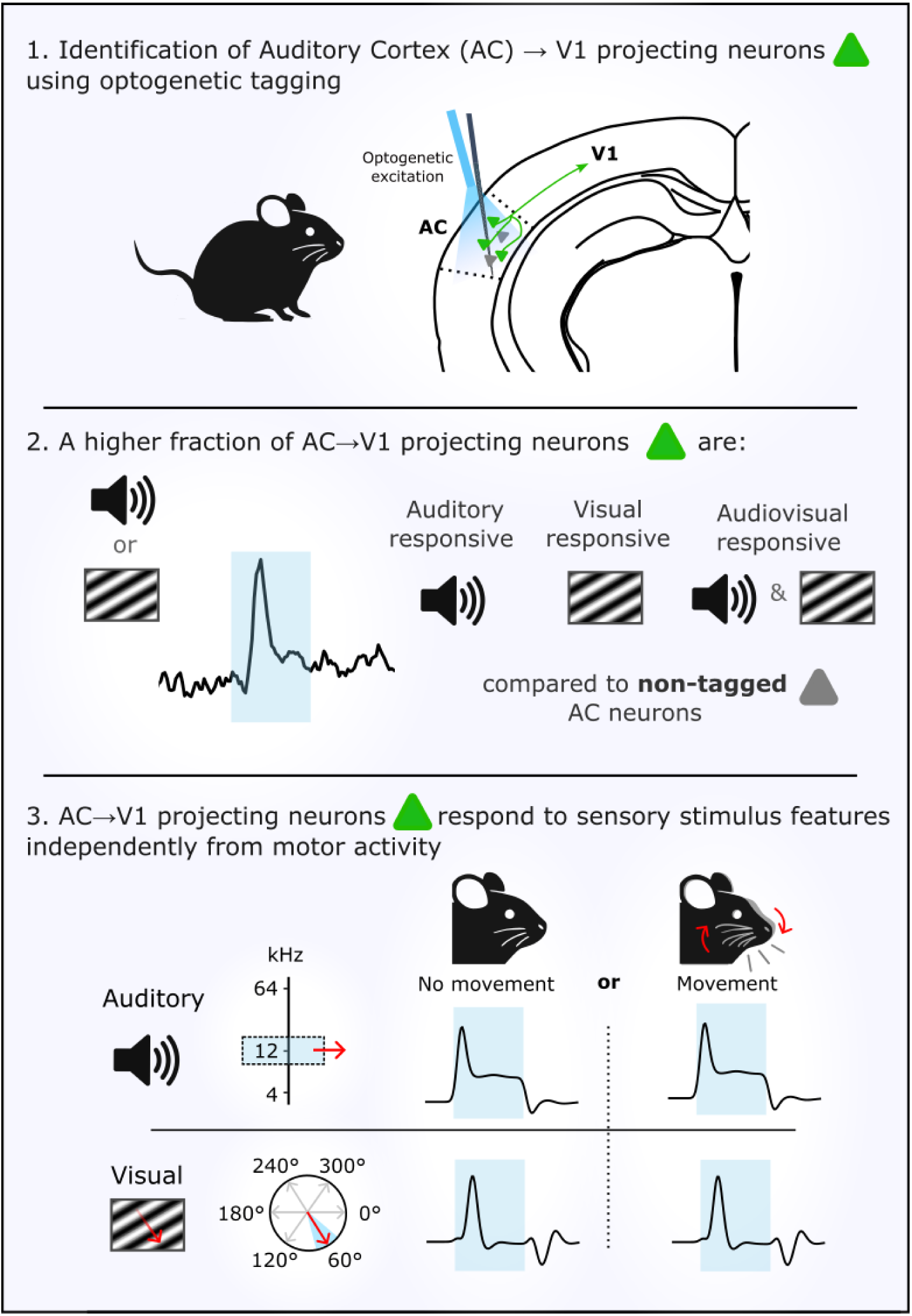

## Introduction

Understanding how the brain integrates information across sensory modalities is a fundamental question in neuroscience. While sensory cortices were classically considered modality-specific, it is now well established that neurons in lower sensory areas respond to stimuli outside their primary modality^1^. For example, cells in mouse primary visual cortex (V1) not only respond to visual stimulation, but are also modulated by sound^2–4^. While part of this sound-evoked modulation in V1 can be explained by the motor behaviour of the animal^5,6^, other components are best explained by auditory but not motor variables^7,8^. Identifying the source and nature of this auditory input to V1 is therefore key to understanding how cross-modal integration is implemented in the cortex.

One possible route runs to V1 via monosynaptic projections directly from auditory cortex (AC). Direct anatomical connections between the rodent AC and V1 have long been established^9,10^. Such connections have also been found in macaques^9,11,12^, suggestive of a conserved evolutionary role. Cross-modal interactions may directly inform processing in the visual cortex, which could help shape visual percepts (e.g. the collision illusion)^13^. Furthermore, conscious perception has been proposed to require widespread multisensory interactions^14–16^. From this viewpoint, the continuous correlation and decorrelation of different sensory streams allow for the assignment of a perceptual interpretation of the then-current activity patterns. Without such cross-modal interactions, the identity of activity in any sensory area remains ambiguous, because there is no other activity pattern to compare it to.

Recent studies have started to explore the information that is transmitted from AC-to-V1 by specifically imaging neurons that are monosynaptically projecting to V1^17–19^. These neurons reside mostly in deeper layers (Layer 5/6) of AC^3,17^, and appear to respond selectively to loud sound onsets when compared to other non-V1-projecting neurons of the AC^17^. When axonal boutons of AC projections to V1 were imaged during a conditioning paradigm, these axon terminals not only responded to auditory, but also to visual stimuli^18^. Consistent with this is a recent finding of a population of visually responsive neurons in layer 5/6 of the AC^20^, raising the possibility that these neurons may overlap with those projecting to V1.

Nonetheless, several important questions remain to be answered. First, although action potentials typically reliably propagate from the soma towards the axonal boutons, their calcium signals may be affected by presynaptic modulation^21^. Thus, to what extent can these axonal bouton results be translated to somatic responses of the cells of origin? Second, and perhaps most importantly, it is unknown whether multisensory response properties are a distinctive feature of AC-to-V1 projecting neurons, or whether they are shared by the broader AC population. This question cannot be resolved by axonal bouton imaging. Third, can part of the previously described sensory-evoked activity in AC-to-V1 projecting neurons be attributed to orofacial motor signals?

Here, we addressed these questions by performing acute, *in vivo* recordings in the AC of the mouse using high-density silicon probes, while specifically identifying the responses of AC-to-V1 projecting neurons by optogenetic tagging. Compared to non-tagged neurons in the AC, we report that AC-to-V1 projecting neurons have a higher probability of responding not only to auditory, but strikingly also to visual and multisensory stimuli. Furthermore, whereas nearly all AC-to-V1 projecting neurons were frequency-tuned, a subset also exhibited selectivity for the orientation of visual gratings. Crucially, these results could not be explained by motor-related activity or by neuron-level properties such as differences in baseline firing rate, cortical depth of the unit along the recording probe, or waveform type. Together, these results highlight a genuine sensory origin of signals transmitted by AC-to-V1 projecting neurons and provide evidence that the high probability of responding to sensory input is a distinguishing feature of this population compared to non-tagged neurons in AC.

## Results

### Identification of V1-projecting cells in auditory cortex

We recorded single-unit activity from the AC of awake mice while they passively observed audiovisual stimuli. To study the firing behaviour of AC neurons projecting to V1, we identified such neurons using optogenetic tagging (*n* = 13 sessions, *n* = 4 mice). To specifically express Channel Rhodopsin in the targeted population of neurons, a combination of a retrograde Cre-recombinase virus injected into V1 and a Cre-dependent channelrhodopsin (ChR2) virus injected into AC was used (Fig. 1B). The retrograde (Cre-recombinase) virus is transported from axon terminals that arrive in V1 back to their cell bodies throughout the brain. Localised injection of (Cre-dependent) AAV-ChR2 in AC ensured specific targeting of V1-projecting neurons. In line with previously published articles, AC-to-V1 projecting neurons (ChR+ neurons; cells labelled in green) were mostly located in L5/L6 of AC^3,17^ (Fig. 1C), with dendrites visibly reaching towards the upper layers of AC.

**Figure 1.**
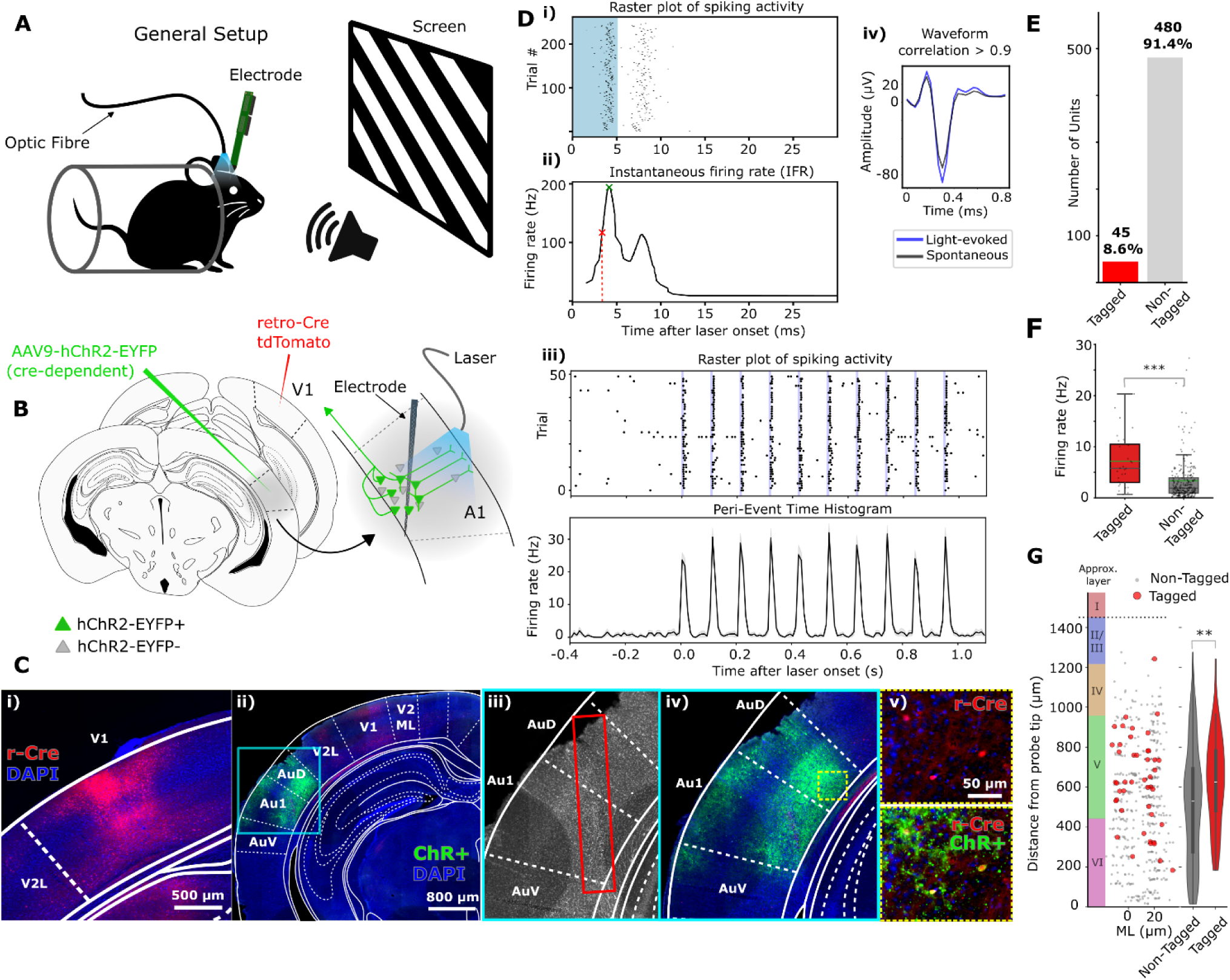
Identification of V1-projecting cells in auditory cortex. **A**) General experimental setup. **B**) Viral injection strategy for targeting V1 projection-specific neurons in the AC. **C**) Example histological images of **i**) virus injection site in V1 (Blue: DAPI, Red: retrograde-Cre (r-Cre) positive tissue), **ii**) Channel Rhodopsin expression in AC (Blue: DAPI, green: Channel Rhodopsin positive), **iii**) Electrode recording track in AC (greyscale, zoomed-in image of (**ii**) using only the DAPI channel). **iv**) Zoomed-in image of light-blue area shown in (**ii**). **v**) Colocalization of retrograde-Cre (Red) and Channel Rhodopsin expression (green). **D**) Units were classified as ‘tagged’ based on four criteria: **i**) Significant ZETA test (p < 0.01, FDR-corrected); **ii**) Laser pulse evoking a firing-rate increase with a spike response latency < 5 ms (defined as first crossing of peak half-height, red cross); **iii**) Reliability of spiking in response to repeated laser pulses in >30% of trials. **iv**) Pearson correlation coefficient between laser-evoked and spontaneous waveforms of > 0.9. **E**) Total number of tagged and non-tagged single units. **F**) Average baseline firing rate tagged versus non-tagged neurons (two-sided Mann-Whitney U test, p = 0.001). **G**) Average distance from probe tip, tagged vs. non-tagged neurons (two-sided Mann-Whitney U test, p = 0.0071). Lateral channel spacing: 20 μm. Colour bar: approximate depth-to-cortical-layer conversion. Abbreviations: V2ML: secondary visual cortex, mediolateral area. V2L: secondary visual cortex, lateral area. Au1: primary auditory cortex, AuV: secondary auditory cortex, ventral area. AuD: secondary auditory cortex, dorsal area.

We targeted AC using 64-channel silicon probes based on stereotaxic coordinates (see *Methods*) and confirmed anatomical placement by robust responsiveness to auditory stimulation at the recording sites. Exact probe placement was further verified using histology (Fig. 1Ciii, Fig. S5). We recorded from a total of 525 single units in the AC, of which 45 (8.6%) were classified as tagged based on their response characteristics to pulsed laser stimulation (Fig. 1D, E, see *Methods*). Tagged neurons showed a significantly higher baseline firing rate compared to non-tagged neurons (two-sided Mann-Whitney U test, p < 0.001) (Fig. 1F). Baseline firing rate was defined as the mean firing rate 500 ms before stimulus onset, averaged over all trials. Additionally, the average location of tagged neurons was slightly more superficial than non-tagged cells (two-sided Mann-Whitney U test, p = 0.0071; Fig. 1G). One explanation for the slight difference in depth could be that the laser light was less likely to reach the bottom-most channels of the probe due to increased light scattering at these depths, resulting in fewer neurons being activated and thus identified as tagged.

### Auditory cortex neurons projecting to V1 are more likely to respond to auditory and visual stimuli compared to non-tagged neurons

To measure the responsiveness of AC-to-V1 projecting neurons to auditory and visual stimulation during electrophysiological recordings, we presented mice with auditory, visual and audiovisual stimuli. Auditory (A) stimuli consisted of single pure tone frequencies ranging from 4-64 kHz; each stimulus was presented at 3 intensities (60, 65, 70 dB). During visual trials (V), one of 6 directional drifting gratings was presented at an identical contrast level (3 orientations). In audiovisual (AV) trials, these auditory and visual stimuli were simultaneously presented. Trial structure was pseudo-randomised for each session, and all stimuli (A, V, AV) were presented for 500 ms, each with an inter-stimulus interval (ISI) of 0.5-1.5 seconds (Fig. 2A). Neurons were classified as auditory or visually responsive using a parameter-free statistical test for detecting time-locked evoked activity (ZETA test, p < 0.05; *see Methods*)^22^. Audiovisual responsiveness required a significant test in both unimodal conditions. Approximately 42.5% (*n =* 223) of the total population of neurons (*n =* 525) recorded in the AC was auditory-responsive. Consistent with the recent literature, we found that about 9.3% (*n =* 49) of the neurons in the AC showed responsiveness to visual stimulation^20^.

**Figure 2.**
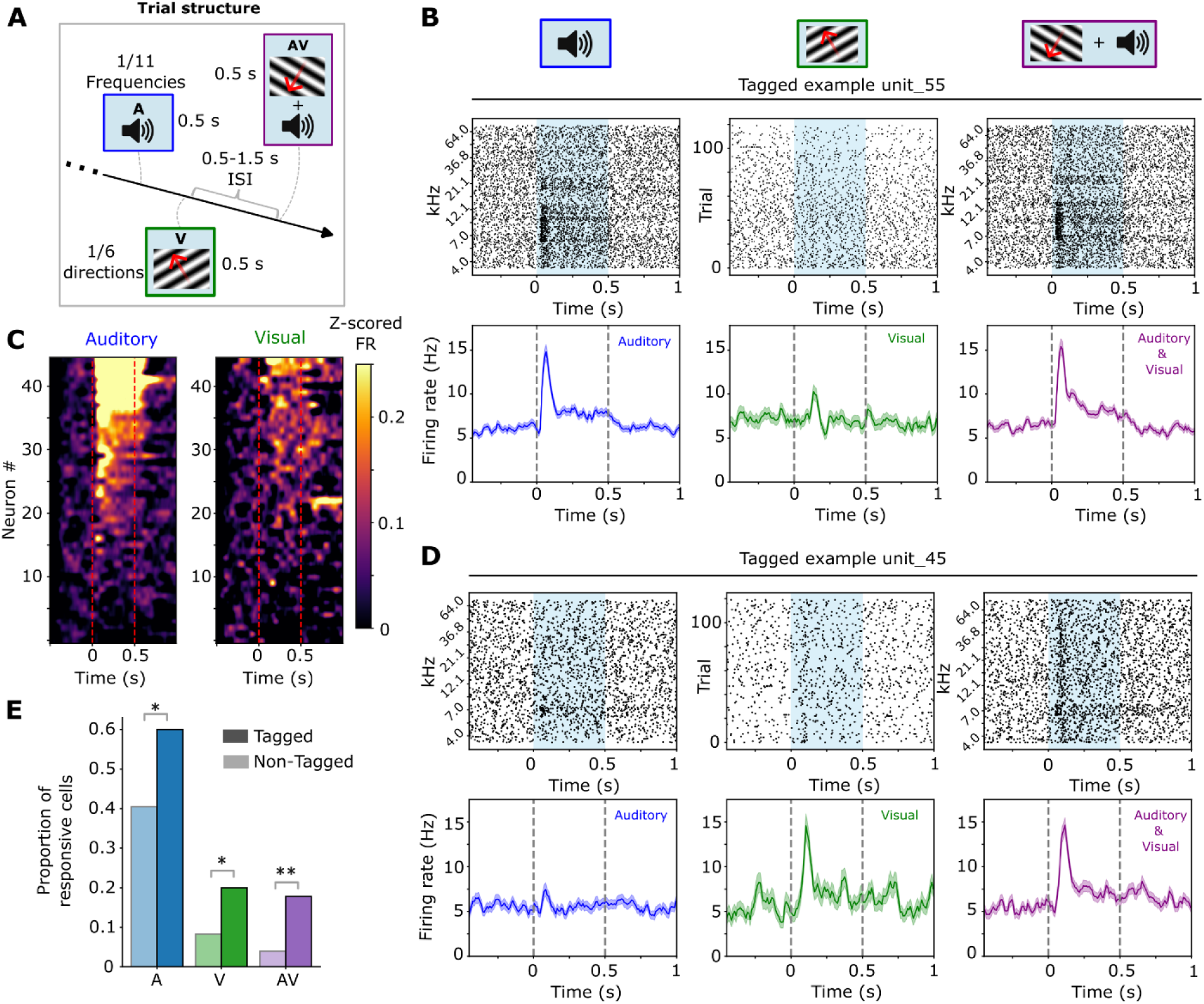
Tagged neurons in auditory cortex are more likely to be auditory, visually and audiovisually responsive compared to non-tagged neurons. **A**) Trial structure. Auditory (A) stimuli (1 of 11 frequencies, 4-64 kHz), visual (V) stimuli (1 of 6 directional drifting gratings) and combined auditory & visual (AV) stimuli were pseudo-randomly presented for 500 ms (inter-stimulus interval (ISI): 0.5-1.5 s). **B**) Raster plots (top) and average firing rate (bottom) of two example tagged neurons during auditory, visual and audiovisual trials, sorted by frequency (auditory & audiovisual plots). Visual trials include all 6 directional drifting gratings (not sorted). Blue: stimulus-on window (0–0.5s). **C**) Average firing rate during auditory (left) and visual (right) trials for all tagged neurons. Individual trials were z-scored (see *Methods*) and averaged over all trials of a specific stimulus condition. Each row represents one neuron, sorted for response magnitude within the stimulus window (0–0.5s). **E**) Proportion of tagged and non-tagged neurons responsive to auditory, visual or both auditory and visual stimuli (two-sided Fisher’s exact test: tagged Auditory vs. non-tagged Auditory, p = 0.017*, Visual, p = 0.026*, Audiovisual, p = 0.001**).

Interestingly, when comparing the AC-to-V1 projecting neurons to the rest of the population, we found that AC-to-V1 projecting neurons were more likely to be auditory, visually and audiovisually responsive (two-sided Fisher’s exact test: tagged Auditory vs. non-tagged Auditory, p = 0.019, Visual, p = 0.017, Audiovisual, p < 0.001) (Fig. 2E). To determine whether audiovisual responses were supra- or sublinear, we calculated the baseline-corrected difference between each unit’s trial-averaged audiovisual response (0-300 ms after stim onset) and the predicted linear summation of both unimodal responses (i.e., response AV – (response(A)+response(V))), for all audiovisually responsive units (ZETA test p < 0.05 for both modalities). For the auditory-only and audiovisual conditions, only trials corresponding to the neuron’s best frequency (BF), defined as the frequency eliciting the highest firing-rate response, were included. The visual response was computed as the average across all visual trials. While the mean response in the audiovisual condition was generally higher than a linear summation of the unimodal conditions in both tagged and non-tagged groups (tagged median difference: 1.30 Hz; non-tagged: 0.34 Hz), indicating a tendency towards supralinearity; however, these differences were not significant (Wilcoxon signed-rank test: tagged, p = 0.41; non-tagged, p = 0.12).

To account for the hierarchical structure of the data and uneven sampling of neurons across animals, we further validated these results using a Bayesian binomial mixed-effects model with random intercepts and slopes per animal. This analysis confirmed that the effect of the ‘tagged status’ (i.e. whether a neuron was identified as tagged) remained positive across auditory (β = 0.874 (log-odds), Odds Ratio (OR) = 2.40), visual (β = 0.888, OR = 2.43), and audiovisual (β = 1.425, OR = 4.15) conditions, with 95% credible intervals excluding zero in all cases. Random slope variances were small across modalities (Auditory (log) Standard Deviation (SD) = 0.197, 95% CI: 0.115–0.338; Visual SD = 0.247, 95% CI: 0.145–0.419; Audiovisual SD = 0.328, 95% CI: 0.195–0.551). These results indicate that the increased sensory responsiveness of AC-to-V1 projecting neurons is robust to uneven sampling across animals and is broadly consistent across individuals.

The median latency in the response to auditory stimuli was approximately 12 ms (IQR: 8-27 ms), generally consistent with expected neural response latencies in the AC to sound^23^. Median response latency to visual stimuli was 85 ms (IQR: 46-120 ms). In V1, latencies to visual stimuli are typically much shorter, around 40 ms^24^. These results are thus consistent with indirect or feedback input from V1, rather than a more direct pathway via the thalamus. Latencies to auditory and visual stimuli were not significantly different between AC-to-V1 projecting neurons and the rest of the population (two-sided Mann-Whitney U test: tagged vs. non-tagged Auditory, p = 0.45, Visual, p = 0.26).

### Auditory cortex neurons projecting to V1 convey feature-specific stimulus information

Neurons in the AC of many species selectively respond to specific sound frequencies. Neurons with similar tuning characteristics cluster together, forming a tonotopic map across the auditory cortical surface^25,26^. Auditory-responsive neurons (*n =* 223) in our recordings showed frequency selectivity (see Fig. 3A for an example), with some neurons being more broadly tuned than others. Furthermore, neurons tended to respond in an intensity-dependent way (Fig. 3A).

**Figure 3.**
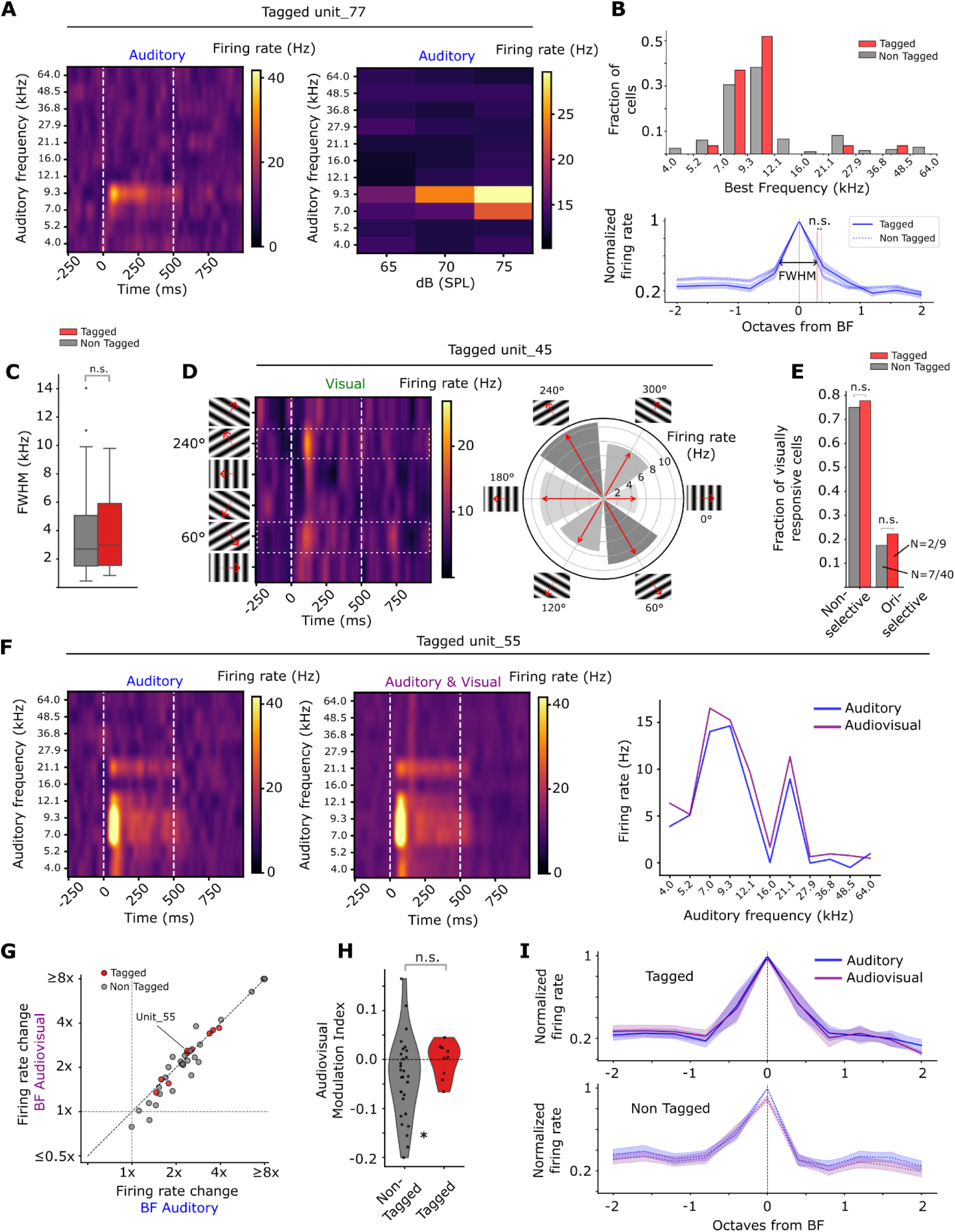
Auditory cortex neurons projecting to V1 convey stimulus feature information to V1. **A)** Left: Firing rate map (Hz) in response to pure tones for an example tagged unit, sorted by frequency. White dashed lines: stimulus on- and offset. Right: frequency-intensity plot for the same unit (tagged unit_77) across stimulus intensities (60, 65, and 70 dB sound pressure level (SPL). **B**) Top: Histogram of best-frequencies (BF) for tagged vs non-tagged auditory responsive AC neurons. Bottom: Comparison of tuning curve width of tagged vs. non-tagged neurons. Tuning curves are normalized and aligned by BF response (mean +/-SEM). **C**) Full-Width-Half-Maximum (FWHM) of auditory tuning curves (Fig. 3B) of tagged vs. non-tagged auditory responsive AC neurons (two-sided Mann-Whitney U test, p = 0.57). **D**) Left: Firing rate map (Hz) in response to visual moving gratings for an example tagged unit sorted by moving grating direction. White dashed boxes signify trials of the same orientation. Note the higher firing rate responses to stimuli with 60° and 240° direction, sharing the same orientation. Right: Average firing rate per visual stimulus direction, rendered in a polar plot; this neuron was found to be significantly orientation selective (OS) (p < 0.05, Hotelling’s-*T*² on individual orientation vectors, see *Methods*). Darker grey: higher firing rate. **E**) Comparison of fraction of visually responsive AC neurons selective for orientation (OS) or non-selective (NS); tagged vs. non-tagged neurons (two-sided Fisher’s exact test: OS, p = 1.00, DS, p = 1.00). **F**) Auditory (left) and audiovisual (middle) firing rate maps (Hz) for an example tagged unit, sorted by auditory frequency. Right: Auditory tuning curves for auditory-only (A) vs. audiovisual (AV) for the same unit (tagged unit_55). **G**) Firing rate (FR) change at BF during stimulus compared to average baseline for all audiovisually responsive AC neurons; A trials (x-axis) and AV trials (y-axis). **H**) Audiovisual modulation Index (MI) per neuron, split into tagged and non-tagged populations (two-sided Mann-Whitney U test, p = 0.12). MI = (FR change AV – FR change A)/(FR change AV + FR change A). Non-tagged cells showed a mean MI significantly below zero (Wilcoxon signed rank test, p = 0.018*). **I**) Comparison of tuning curves during auditory vs. audiovisual trials for tagged (top) and non-tagged (bottom) audiovisually responsive neurons. Tuning curves are normalized and aligned by BF response (mean +/-SEM).

To test whether V1-projecting AC neurons showed differences in frequency selectivity relative to non-tagged neurons, we compared the distributions of best-frequencies (BF) between the two populations^23^. Most neurons in our recordings had BFs in the sub-ultrasound range (approximately 6-12 kHz), with a few neurons sensitive to lower and higher frequencies (Fig. 3B). The distribution of best frequency values did not differ significantly between V1-projecting and non-tagged neurons (Kolmogorov-Smirnov test, p = 0.60), suggesting no systematic difference in frequency preference between the two populations. Next, we calculated the full width at half-maximum (FWHM) of each tuning curve to compare curve widths between neurons (see *Methods*). The mean FWHM did not differ significantly between tagged and non-tagged cells (two-sided Mann-Whitney U test, p = 0.57; Fig. 3B-C).

Having characterised auditory tuning, we next wondered whether visually responsive AC neurons showed any selectivity to visual features such as orientation and direction of moving gratings. We quantified the degree of orientation and direction tuning using a previously published vector-based approach^27^. A modest fraction of all visually responsive AC neurons showed significant orientation selectivity (9/49, Hotelling’s T² on individual orientation vectors, p < 0.05), a proportion significantly exceeding chance (binomial test, p < 0.001). Notably, a small subset of orientation-selective neurons were tagged AC-to-V1 projection neurons (2/9, Fig. 3D, E), suggesting that the AC does not only convey visual signals to V1 in a broad sense, but also transmits feature-specific visual information. Additionally, some individual neurons were found to be direction-selective (n = 3/49, Student’s T-test, p < 0.05, see *Methods*); however, this fraction could be expected by chance alone (binomial test, p = 0.45). No differences were observed in the fraction of orientation-selective neurons between tagged and non-tagged populations (Fig. 3E).

To investigate whether concurrent auditory and visual stimulation modulates auditory tuning properties, we compared audiovisually responsive neurons during auditory-only (A) and audiovisual (AV) trials. Specifically, we compared for each neuron the average evoked firing rate at its best-frequency (BF) during auditory-only trials with the firing rate at that same BF during audiovisual trials. At the single-neuron level, none of the audiovisually responsive neurons significantly changed their average firing rate at BF in audiovisual compared to auditory-only trials (two-sided Mann-Whitney U test, p < 0.05; Fig. 3G). To assess group-level differences in audiovisual modulation, we computed an audiovisual modulation index ((FR change BF(AV) – FR change BF(A)) / (FR change BF(AV) + FR change BF(A))). Non-tagged neurons showed a small but significant suppression of responses to audiovisual stimuli relative to auditory alone, as reflected by the modulation index being significantly below zero (Wilcoxon signed-rank test, p = 0.018, Fig. 3H). Tagged neurons, in contrast, showed no consistent modulation (p = 1.0). No significant difference was found between the two groups (Mann-Whitney U test, two-sided, p = 0.122, Fig. 3H). Besides changes in BF, neurons could also shift their tuning curve due to the addition of visual stimulation. To test this possibility, we compared the average tuning curves of audiovisually responsive neurons for both populations during auditory-only trials and audiovisual trials. The shape of the audiovisual tuning curves for both populations (tagged and non-tagged) during auditory-only and audiovisual trials largely overlapped (Fig. 3I), indicating that the addition of visual activity does not significantly change the selectivity to auditory stimuli (FWHM auditory-only vs. FWHM audiovisual; Wilcoxon signed-rank test, tagged; p = 0.43, non-tagged; p = 0.33).

Furthermore, while the overall orientation selectivity slightly decreased in the AV condition (normalised length of the mean response vector; Wilcoxon signed-rank test, p = 0.039, see *Methods*), the majority of orientation-selective neurons remained significantly tuned in the AV condition (6/9, Hotelling’s T², p < 0.05). Taken together, these results suggest that combined audiovisual stimulation leaves tuning properties of AC neurons largely intact.

### Disentangling sensory and motor-related contributions to auditory and visual responses in auditory cortex

Movement is known to influence cortical activity throughout the brain^28–30^. Therefore, we sought to examine whether motor movement could explain part of the stimulus-evoked firing patterns we observed. We quantified the amount of motor activity during each session using FaceMap^31^ and used the first ten principal components (PCs) obtained from the absolute frame-to-frame differences to quantify the amount of motion energy (ME) during individual trials. We calculated ME for both orofacial movements using a video of the face of the mouse, and for body movements using a separate camera positioned to capture large movements of the trunk and limbs (Fig. 4A, Fig. S1A-D; see *Methods*). Subsequently, we split all trials into two groups based on the amount of ME present (Fig. S2A). Trials that exceeded 2 SD of the ME distribution during stimulus presentation were classified as “high movement trials”; all other trials with little to no movement were classified as “low movement trials”.

**Figure 4.**
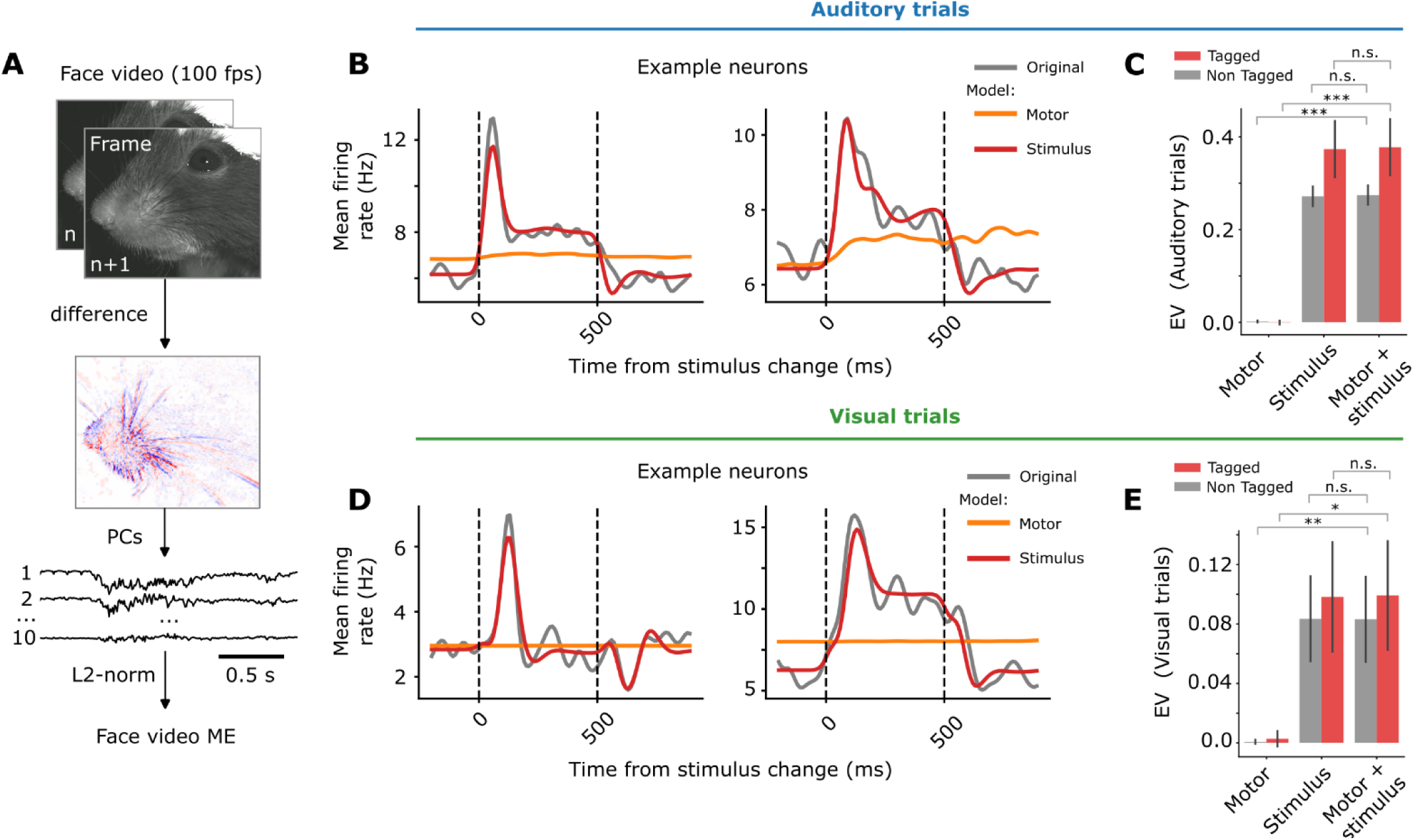
Motor-related activity contributes minimally to the average firing rate of tagged and non-tagged auditory and visually responsive neurons in auditory cortex during sensory stimulation. **A)** Orofacial motion energy (ME) was defined as the absolute sum of squares of the first 10 principal components (PCs) of the frame-to-frame differences of the face video. Middle: red/blue colour: increased/decreased pixel intensity. **B)** Left: Average single-neuron firing rate of auditory-responsive neurons was predicted by a generalised linear model (GLM) using motor predictors (orange), stimulus predictors (red), or both predictors (not shown). Dotted lines: stimulus on- and offset. Two example model fits are shown for two AC neurons. **C)** Cross-validated explained variance (EV) in mean firing rate (0-1s relative to stimulus onset) during auditory trials, scored separately for each individual model (Wilcoxon signed-rank test motor-only model vs. motor & stimulus model; tagged neurons: p < 0.001***, non-tagged neurons: p < 0.001***. Stimulus-only model vs. stimulus & motor model; tagged neurons: p = 0.09, non-tagged neurons: p = 0.20. **D**) Same as **B**), for visually responsive neurons. **E**) Same as **C**), for visual trials (Wilcoxon signed-rank test motor-only vs. motor & stimulus; tagged: p = 0.039*, non-tagged: p < 0.01**. Stimulus-only model vs. stimulus & motor model; tagged: p = 0.82, non-tagged: p = 0.50.

Auditory stimuli were generally more likely than visual stimuli to evoke movements of the facial musculature of the mouse (Fig. S1E, F); on average, the ME on 11.8 ± 1.5% of all auditory trials exceeded 2 SD, contrasted to 8.0 ± 1.1% during visual trials (Wilcoxon signed-rank test, p = 0.047). We found similar results when analysing body movements of the mouse (15.9 ± 1.3% of auditory trials exceeded 2 SD; 12.6 ± 1.4% of visual trials, Wilcoxon signed-rank test, p = 0.032). These results are in line with previous literature^8,32^, although the proportion of trials with movement during auditory stimulation is much larger in the paper by Olsen and Hasenstaub^32^ (61% vs.11.8% in this study), possibly due to differences in auditory stimulation type (noise bursts versus pure tones in this study).

We next compared the average neuronal firing rates during auditory trials with high versus low ME for each neuron individually. A large number of AC neurons were significantly associated with face motion energy, as their firing rates increased significantly during trials with high ME (98/223 = 43.9% of neurons, Fig. S2C, D, left panel). The number of neurons that significantly changed firing rate on high ME compared to low ME during visual trials was much lower (10/49 = 10.2% of neurons; Fig. S2E, F; two-sided Fisher’s exact test: p < 0.001). The proportion of AC-to-V1 projecting neurons that significantly changed firing rate on high vs. low ME trials was not different compared to non-tagged neurons (Fig. S2D, F; neither in auditory nor visual trials).

While AC firing rates were associated with motor activity in a substantial proportion of neurons during auditory trials and to a lesser degree during visual trials, it remains unclear to what degree motor activity constitutes a primary factor driving these firing rate responses. We therefore sought to further disentangle sensory-driven activity from motor-related contributions to the average firing rate of AC neurons. We employed a time-resolved generalized linear model (GLM) to predict single-neuron activity based on two sets of predictors: motor-related and stimulus-related predictors. We fit and compared three models per neuron: one using only motor predictors, one using only stimulus-related predictors and one using both motor predictors and stimulus predictors (see *Methods*). Averaging the predicted firing rate across trials revealed that motor activity contributed minimally and only sporadically to single-neuron responses in AC in both auditory and visual trials (Fig. 4B, D).

To quantify the contributions of motor-related and sensory signals to stimulus-evoked responses, we scored the predicted firing rates of each GLM model against the actual average firing rate using five-fold cross-validated explained variance (EV; see *Methods*); each model of each neuron was evaluated individually. This analysis showed that, for both modalities, the average time-varying firing rate in AC was best explained by auditory and visual predictors, whereas motor predictors explained little to no variance (Fig. 4C, E). Indeed, using both motor and stimulus predictors significantly improved explained variance compared to motor-only models, in both tagged and non-tagged populations (Wilcoxon signed-rank test: motor-only model vs. motor & stimulus model; tagged neurons: p < 0.001, non-tagged neurons: p < 0.001). Furthermore, adding motor predictors to the stimulus-only model did not significantly increase explained variance in both unimodal conditions (Fig. 4C, E). This indicates that motor movements are not consistently driving stimulus-locked neural activity in these neurons. Taken together, we conclude that, while motor movement may modulate AC firing rates, sensory signals constitute the primary drive of the stimulus-evoked responses identified in this study.

### Classification of broad-and narrow-spiking neurons

Next, we characterised the tagged and non-tagged populations by examining their spike waveform properties, as this could further reveal their cell-type identity (i.e. excitatory versus inhibitory). We analysed a subset of sessions (*n =* 6/13 sessions, *n* = 293 units), as slight differences in filter settings in some sessions precluded reliable comparison of waveform shapes in the full dataset. We classified neurons as broad-spiking (BS) and narrow-spiking (NS) using the peak-to-trough ratio and peak-to-trough delay of the waveform (Fig. 5A). Narrow-spiking neurons are typically classified as interneurons, whereas broad-spiking neurons are regarded as excitatory neurons^33^. Consistent with known proportions of interneurons and excitatory neurons in the mouse cortex, about 15% of the population of neurons recorded in our dataset was classified as NS, whereas 84.7% was classified as BS, and 0.3% were non-classifiable^34^. About 31% of tagged neurons were classified as NS, compared to 69% classified as BS (total *n =* 29 tagged units; Fig. 5A). This finding was surprising, given that projection neurons are most frequently glutamatergic pyramidal neurons^35^, although there is evidence that extra-areal projection neurons can be inhibitory^36,37^.

**Figure 5.**
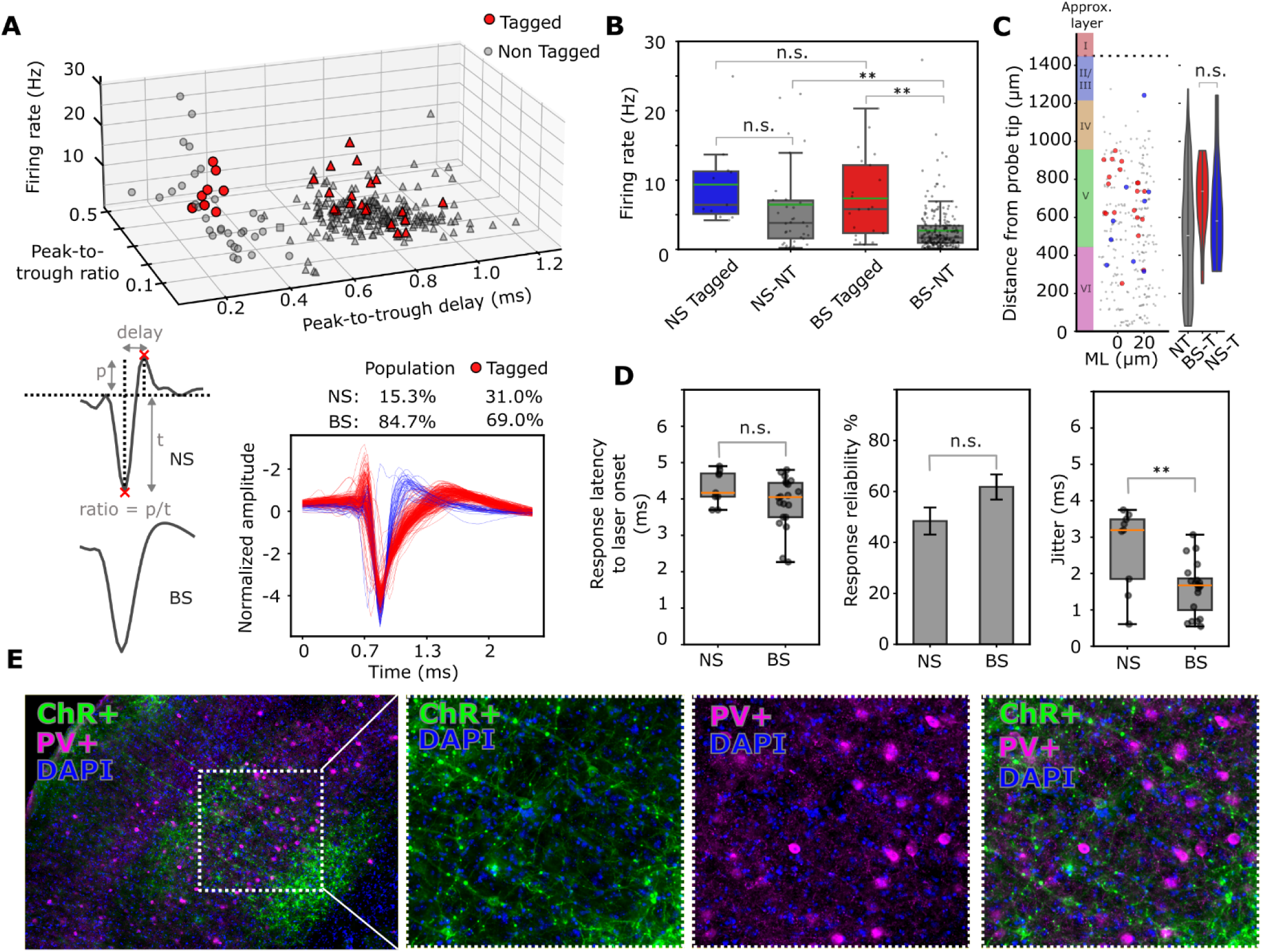
Classification of broad- and narrow-spiking neurons. **A**) Top: Neurons were classified as putative inhibitory, narrow-spiking (circles) or excitatory, broad-spiking cells (triangles), or non-classifiable (squares) using two criteria: 1) Peak-to-Trough delay in ms, 2) Peak-to- trough ratio. Red: tagged neurons. Bottom left: Classification criteria visualised (p = peak, t = trough). Bottom right: waveforms for neurons classified as broad-(red) and narrow-spiking (blue). **B**) Baseline firing rate comparison between tagged vs. non-tagged (NT), broad-(BS) and narrow-spiking (NS) units (NS-tagged vs. NS-NT, p = 0.08, NS-tagged vs. BS-tagged, p = 0.35, NS-NT vs. BS-NT, p < 0.001**, BS- tagged vs. BS-NT, p < 0.001**, two-sided Mann-Whitney U test). **C**) Recording depth of broad- and narrow-spiking tagged neurons (two-sided Mann-Whitney U test, p = 0.15; ML: mediolateral, NT: non-tagged, NS-T: narrow-spiking tagged, BS-T: broad-spiking tagged). Colour bar: approximate depth-to-cortical-layer conversion. **D**) Left: spike response latency to laser onset (one-sided Mann-Whitney U test, p = 0.072). Middle: reliability of response to laser pulse (one-sided Mann-Whitney U test, p = 0.075). Right: jitter in first spike latency relative to laser onset (one-sided Mann-Whitney U test, p = 0.0075). Orange line: median response. **E**) Left: example histological image of Channel Rhodopsin-positive neurons (ChR+, green), parvalbumin-positive neurons (PV+, purple) and DAPI (Blue). Right panels: No overlap was observed between ChR+ and PV+ neurons (*n* = 2 animals, 4 sections per animal).

However, given the relatively high proportion of NS units among tagged neurons, indirect activation may provide an alternative explanation. Fast-spiking interneurons are tightly coupled to local pyramidal neurons and may be indirectly activated by photostimulation of pyramidal neurons, even at short latencies^38^. Therefore, we first asked whether the NS- and BS-tagged groups differed in the parameters used as tagging criteria. We found no significant differences between NS- and BS-tagged neurons in these parameters (firing-rate response latency to laser onset, NS-tagged vs BS-tagged: one-sided Mann-Whitney U test, p = 0.072; reliability of spiking to laser pulse, NS-tagged vs. BS-tagged: one-sided Mann-Whitney U test, p = 0.075; Fig. 5D: left, middle panel), arguing against indirect activation of these tagged NS neurons. To further investigate the possibility of indirect laser-induced activation of tagged neurons classified as NS, we compared the standard deviation of the first-spike latency following laser onset (commonly referred to as ‘jitter’) between NS- and BS-tagged neurons. The population of tagged NS neurons showed, on average, slightly higher jitter values compared to BS-tagged neurons (one-sided Mann-Whitney U test, p = 0.0075). However, considerable overlap was observed, suggesting that jitter alone cannot reliably distinguish between directly and indirectly activated neurons, consistent with previous reports^39^.

To further examine whether tagged NS neurons could be indirectly activated, we estimated the probability of monosynaptic connections by computing spike time cross-correlograms (CCGs) between all tagged neuron pairs (Fig. S3A). If many of our NS-tagged neurons were indirectly activated, they were predicted to show a high probability of being monosynaptically connected with other tagged neurons in our population. Neurons were considered monosynaptically connected to an excitatory cell if the CCGs showed a short, positive latency peak within 1-4 ms, larger than the global 99% band of spike counts^40,41^ (see *Methods*). Although the proportion of monosynaptic connections between all NS and BS pairs was slightly higher between tagged neuron pairs (2/40, 5%) compared to the non-tagged group (32/1269, 2.52%), this difference was not significant (two-sided Fisher’s exact test, p = 0.28; Fig. S3D). Taken together, while we cannot fully exclude the possibility of some of our units being indirectly activated, the broader pattern of results does not support indirect activation as a predominant explanation, thus favoring the possibility of a population of NS neurons in AC that project to V1.

To further characterise the genetic identity of these NS-tagged neurons, we asked whether they might correspond to parvalbumin-positive (PV+) interneurons, which account for the majority of neocortical interneurons^42^ and include fast-spiking basket cells and chandelier cells^43,44^. We therefore performed an additional secondary antibody staining for PV+ (Fig. 5E). No overlap between ChR+ neurons and PV+ labelling was found, indicating that the population of NS-tagged neurons did not express parvalbumin.

### Basic neuronal properties cannot explain sensory response differences between V1-projecting and non-tagged neurons

The observed differences in sensory responsiveness between AC-to-V1 projecting neurons (tagged) and non-tagged neurons could potentially reflect underlying differences in neuronal properties, such as motor correlates, baseline firing rate or recording depth, rather than a genuine difference in sensory processing. To rule out this possibility, we fit a logistic regression model with stimulus responsiveness (ZETA test, p < 0.05) as the outcome variable. In addition to a binary variable indicating whether a neuron was significantly associated with motor activity (Fig. S2C, E), we tested whether recording depth and baseline firing rate correlated significantly to a neuron’s sensory responsiveness (see *Methods*).

This analysis revealed that AC-to-V1 projecting neurons were significantly more likely to be auditory-responsive than non-tagged cells, even after controlling for motor activity, depth, and spontaneous firing rate (Odds Ratio (OR) = 2.9, 95% CI [1.46, 5.56], p = 0.002). As expected, motor activity was significantly associated with auditory responsiveness (OR = 3.9, 95% CI [2.60, 5.89], p < 0.001; cf. Fig. S2), while recording depth showed a small but significant negative effect (OR = 0.999, p = 0.010), indicating that neurons located slightly more superficially to the cortical surface were somewhat less likely to be auditory responsive. Critically, after controlling for motor modulation, depth and spontaneous firing rate, AC-to-V1 projecting neurons were still significantly more likely to be visually responsive compared to non-V1 projecting neurons (OR = 2.64, 95% CI [1.16, 6.02], p = 0.021). Baseline firing rate, recording depth and motor modulation were not significantly associated with visual responsiveness.

In a subset of the data (*n =* 6/13 sessions, *n* = 293 units), we reran the logistic regression analysis, further extending the model with a predictor indicating whether a neuron was broad- or narrow-spiking. For both logistic regression models (auditory and visual modalities), adding this predictor did not improve the model, and was not significantly associated with the outcome (p = 0.94, auditory; p = 0.57, visual). We conclude that AC-to-V1 projecting neurons are more likely to be auditory and visually responsive compared to non-tagged neurons, independent of differences in neuronal properties, such as motor correlates, baseline firing rate and recording depth.

## Discussion

We recorded single-unit activity from the AC of awake mice during presentation of audiovisual stimuli while specifically identifying AC neurons projecting to V1 using optogenetic tagging. Compared with non-tagged neurons in our recordings, AC-to-V1 neurons exhibit distinct audiovisual response properties: they are more likely to respond to pure-tone auditory frequencies and to directional drifting gratings. Furthermore, AC-to-V1 neurons were also more likely to respond to combined auditory and visual stimuli. Investigating their responses to different types of auditory stimuli revealed that most AC-to-V1 projecting neurons carried information about the specific frequency presented. Interestingly, some AC-to-V1 projecting neurons also increased their firing selectively to some orientations, but not to others, suggesting a degree of orientation tuning. This indicates that these neurons can carry information about the type of visual stimulus presented. These tuning characteristics were not exclusive to tagged AC neurons. Importantly, these results could not be explained by motor-related activity or differences in baseline firing rate, recording depth and waveform type. Together, these results highlight a genuine sensory origin of information transmitted by AC-to-V1 projecting neurons.

### Visual responsiveness of V1-projecting auditory cortex neurons

A previous study showed that the AC harbours neurons responding to visual stimuli^20^, but did not examine their projection targets. Nonetheless, in line with our results, there is evidence that axonal boutons originating from AC neurons arriving in V1 are visually responsive^18,19^, although the two studies report different response magnitudes. One discrepancy between these studies is the difference in visual stimulus types being used (sinusoidal grating in Garner and Keller^18^, vs. LED flashes in Mazo et al.^19^), while both stimulus types are known to evoke visual responses in AC^20^. Moreover, Garner and Keller^18^ repeatedly paired auditory and visual stimuli with reward across recording days, whereas Mazo et al.^19^ did not; this likely contributed to the observed differences, as visual responses increased across conditioning days in the former study^18^.

Importantly, both Garner and Keller and Mazo et al. measured visually evoked activity of axonal boutons, leaving open the question of whether the somata of AC-to-V1 projection neurons are themselves visually responsive, or whether visual signals emerge through local presynaptic modulation at the bouton level. Although action potentials typically propagate reliably from the soma towards the axonal boutons, presynaptic modulation can take place^21^. In this light, it is interesting that we do find visually responsive AC-to-V1 projection neurons using extracellular electrophysiology, as this method predominantly records spiking activity generated near the soma and axon initial segment^45,46^.

Our observations are consistent with the notion that AC-to-V1 projecting cells convey visual information to V1, especially because a small but significant subset of AC-to-V1 neurons did so in an orientation-selective manner (Fig. 3D, E). However, visually responsive neurons in AC were previously not found to be orientation-selective^20^. One possible explanation for the difference is the smaller number of visually responsive neurons recorded (*n =* 0/7) in the previous study by Morrill and Hasenstaub^20^, which may have been insufficient to detect significantly orientation-tuned cells, also given the rather low probability of significant orientation tuning we found (Fig. 3E).

### Motor correlates during sensory-evoked activity in auditory cortex

Over the years, it has become increasingly clear that motor movements heavily influence firing rates across the brain^28–30^. Auditory stimuli can elicit time-locked stereotyped movements^47^, making it essential to control for multisensory experiments^48^. Our results confirm that face movements were present in a substantial fraction of auditory and visual trials. A comparison of average firing rates during auditory trials with high facial or body movements with low-movement trials revealed that a substantial fraction (43.9%) of auditory-responsive neurons increased their firing rate on these high-movement trials (Fig. S2).

We did not expect to see that such a large fraction of neurons would be significantly associated with movement, as other studies reported substantially smaller fractions of neurons in AC. Clayton et al. report that about 11% of the neurons in L2/3 and L4 were significantly modulated by movement, whereas up to 25% of the regular spiking (RS) neurons in L5/6 of AC changed their firing rates directly following spontaneous facial movements^47^. However, when we separated motor-related neural responses from sensory-evoked responses in AC using a GLM approach, we found that the average evoked firing rate of individual sensory-responsive neurons could not be explained well by movement alone.

Part of this difference in the fraction of movement-modulated neurons could result from a selection bias; the selection of high-motion-energy trials may have unintentionally favored specific auditory frequencies that themselves drove the stronger sensory responses in these AC neurons, rather than movement being the underlying factor. Nevertheless, pure tone frequencies at 60, 70, or 75 dB are not thought to reliably elicit stereotypical movements in mice^47^, suggesting that auditory and motor contributions are unlikely to be systematically confounded in the current dataset, allowing them to be separated. While motor activity may still contribute to a general increase or decrease in firing rate in a subset of the population, the GLM results indicate it cannot account for the auditory responses of AC neurons, suggesting that sensory responsiveness remains the primary underlying determinant for the results described in this study.

Previous work demonstrated that some of the auditory-evoked responses in V1 are motor-independent^7,8,19^, contrasting with findings that attribute all auditory-evoked responses in V1 to a motor origin^5^. In addition, it has been reported that AC-to-V1 projection neurons transmit auditory information to V1, with a particular selectivity for loud sound onsets^17^, indicating that motor-independent auditory signals in V1 could depend on AC-to-V1 projection neurons. However, these latter conclusions were primarily based on broadband, white-noise auditory stimuli, which reliably evoke stereotypical movements even at moderate intensity levels^47^. As mentioned, neurons in different layers of auditory cortex could be differentially modulated by motor movements, with neurons in deeper layers being affected more prevalently^49,50^, precisely where AC-to-V1 projection neurons tend to be situated^3,17^. Consequently, it remains difficult to ascertain whether, and to what extent, the results reported by Deneux et al.^17^ can be attributed to sensory versus motor processing. Our results help to resolve this ambiguity, providing evidence that AC-to-V1 projection neurons can convey auditory and visual information, and not only motor correlates. Our findings indicate that AC-to-V1 projection neurons directly convey motor-independent signals to V1, although we did not directly measure their effects on V1. Yet our results do not rule out the possibility that a subset of AC-to-V1 projection neurons primarily convey motor-related activity.

### Spike-waveform diversity among V1-projecting auditory cortex neurons

A relatively large fraction of our tagged neurons was classified as narrow-spiking (31% vs. 15% of the total recorded population). Narrow-spiking neurons are commonly considered interneurons^33^, although extracellular spike width alone does not provide a definitive classification, as there is some evidence for pyramidal neurons showing “thin spikes” when identified via antidromic pyramidal tract stimulation^51^. Nonetheless, this raises the possibility that a subset of AC neurons classified as tagged correspond to GABAergic interneurons. This finding was unexpected, as most neurons that project to distant parts of the cortex originate from pyramidal neurons^35^. Nevertheless, there is accumulating evidence suggesting the existence of GABAergic interneurons projecting to distant cortical and subcortical targets in the neocortex^36,37,52,53^, with most corticocortical GABAergic neurons being somatostatin-positive^52^. While there is no clear consensus regarding the function of long-range inhibitory connectivity, one function could be the temporal coordination of activity in distant brain regions^54^. GABAergic interneurons are considered especially fit for this purpose because of their extensive axonal arborisation within target areas, as well as their highly myelinated axons, which facilitate fast action potential propagation^37^.

### Functional significance of audiovisual cross-modal interactions

Our results indicate that AC-to-V1 projection neurons convey both auditory and visual information to V1 with a higher probability than found in the general population of AC neurons. What could be the functional significance of these cross-modal signals? What computational role may be served by informing the visual cortex about auditory and, especially, visual information emitted by auditory cortex? One explanation might be offered by predictive coding theory^18,55–58^. Under this view, AC-to-V1 projection neurons could predict the current visual input, either directly by way of visually responsive AC cells or indirectly, through acquired associations between certain auditory and visual stimuli. This view is supported by evidence showing that learned associations between auditory and visual stimuli lead to experience-dependent suppression of visual responses of axonal boutons in V1^18^. In this theoretical framework, transmission of visual information from AC-to-V1 would reflect a best-guess representation of which sensory input is most likely to reach V1.

A second, complementary mechanism may involve conditional gating of sensory information via recurrent feedback loops. Here, the input to AC from one modality, e.g. visual information to AC, modulates the gain of AC output, selectively enhancing or suppressing the transmission of sensory signals when they align well and coincide with other inputs to AC (e.g. from somatosensory or motor areas^59^), setting the context in which visual information is embedded. More broadly, the genesis of a percept, be it auditory or visual, should not be seen as a standalone process occurring in one particular area, but can be understood as a generative result of the interaction with other modalities^59^. This theoretical perspective suggests that computing correlations and de-correlations across large, multisensory networks may serve both the integration and segregation of sensory modalities^14,60^. We hypothesise that AC-to-V1 neurons may play an active role in this multisensory cortical circuit, potentially contributing to predictive and context-dependent modulation of visual processing through cross-modal feedback and gain control mechanisms.

### Limitations of the study

First of all, we cannot fully exclude the possibility that a fraction of neurons labelled as tagged was actually indirectly activated by light. However, we applied several criteria to minimise this potential risk, thus ensuring that the population of tagged neurons should reflect to a very large extent AC-to-V1 projection neurons. Whether or not narrow-spiking neurons can be identified as GABAergic interneurons or interareally projecting neurons remains to be investigated in future studies. Third, the total number of directional drifting gratings presented in our study was limited, constraining the resolution at which tuning curves could be estimated. Further experiments with denser sampling of stimulus orientations are required to characterise their tuning properties more precisely.

## Methods

### Animals

All experiments were approved by the Dutch Committee for Animal Experiments and by the local Animal Welfare body of the University of Amsterdam. A total of six male wild-type (C57BL/6, Envigo) mice were used in this study. Four animals were used in the electrophysiological experiments, and two animals were used for immunohistological experiments to test for overlap between PV+ and ChR+ neurons (Fig. 5E). Mice were at least eight weeks of age at the start of the experiments. Animals were housed in pairs of two and kept on a reversed day-night schedule (lights-off 8:00-20:00 hrs), in temperature-controlled rooms (21± 2° C, humidity 55% ± 15%). Individual home-cages were divided into two equally large, separated compartments to prevent the possibility of fighting.

The overall chronology of the experiment was as follows: (1) animals were habituated to the experimenter for at least five days; (2) head-bar and viral injection surgery were combined on a single day; recovery took at least 2 days; (3) Mice were habituated to head-fixation in the experimental setup for at least five days for up to 10 days, increasing the total duration of head-fixation daily until approximately 1 hour; (4) Craniotomy surgery was executed; recovery took at least 24 hours; (5) Electrophysiological data were recorded daily for a maximum of seven days; (6) Animals were perfused at the end of the experiment.

### Surgical procedures

#### Head-bar implantation

Mice were anaesthetised using isoflurane (4-5% for induction, ∼2 % maintenance) and fixed in a stereotaxic apparatus. A small circular patch of skin covering the skull was shaved off and then cut using scissors, exposing the skull. A titanium head-bar was briefly fixed to the area of exposed skull using super-glue and then properly attached using dental cement (Sun Medical CCB Superbond). Regular dental cement (Paladur) was used to seal the edges between the head-bar and skin. The dental cement was allowed to dry before continuing with virus injections. The head-bar was placed slightly laterally towards the left hemisphere to obtain sufficient space for further procedures.

#### Virus injection

Small holes (∼100 µm) were made in the skull using a dental drill over A1 (relative to bregma: -2.6 AP, 4.3 ML) and V1 (relative to bregma: -3.5 AP, 2.6 ML). Both viral injection locations in A1 and V1 were approached using a straight angle relative to the surgery table. This resulted in a non-perpendicular angle of entry relative to the cortical surface. A total of 138 nL was injected into A1 at a depth of 800 µm in 10 individual steps of 13.8 nL with 1 minute in between injections (pAAV-EF1a-double floxed-hChR2(H134R)-EYFP-WPRE-HGHpA, AAV9; buffer: PBS + 0.001% Poloxamer 188, *Addgene*), further diluted 10 times with sterile saline. The glass pipette was retracted after 15 minutes following the last injection, allowing the virus to diffuse into the tissue. The same amount (138 nL) of retrograde virus was then injected into the craniotomy made over V1 (pAAV-hSyn-Cre-P2A-dTomato, AAV Retrograde; buffer: PBS; 0.001% Poloxamer 188; 200 mM NaCl, Addgene) at two depths, 600µm and 200µm below the cortical surface. At each depth in V1, five individual portions of 13.8 nL were injected with 1 minute in between injections. An interval of at least 10 minutes was used between injections at the different cortical depths. The pipette was retracted after 15 minutes following the last injection. Injections were done with a Nano Inject system (Drummond Scientific) in combination with custom glass pulled pipettes. The skull was covered using two-component silicon putty (Picodent twinsil), over which a hand-cut aluminium cover was glued. Animals were allowed to recover for at least 2 days before the phase of becoming habituated to the experimental setup.

### Stimuli

#### Auditory stimuli

Auditory stimuli consisted of 11 pure tones (sine waves) spaced logarithmically between 4 and 64 kHz (exact stimuli used were: 4000, 5278, 6964, 9190, 12126, 16000, 21112, 27858, 36758, 48503, 64000 Hz). Three different intensity levels were used (65, 70, 75 dB sound pressure level, SPL). Auditory stimuli were presented using a single ES1/ED1 electrostatic loudspeaker/driver (Tucker-Davis) in combination with a Steinberg UR12 external audio card (sampling rate: 192 kHz). During auditory-only trials, one out of 33 frequency-intensity combinations was selected randomly and presented for 500 ms. Each frequency-intensity level was presented 10 times (total number of trials: 330). The speaker was positioned slightly on the right side of the mouse, approximately 10 cm from the head of the mouse. The speaker was calibrated for each frequency-intensity level within 2/3 dB sound pressure level (SPL). Briefly, calibration of the individual frequency-intensity pairs was performed using a microphone sensitive to ultrasound with a known frequency-response curve (Dodotronic, Ultramic 382k EVO). The relative amplitude of each frequency-intensity pair was modified accordingly (increased/decreased) by estimating the actual volume of the speaker, taking into account the frequency-response curve of the microphone at each frequency level.

#### Visual stimuli

Visual stimuli were presented on a 24-inch (30 by 53.5 cm) Dell monitor (resolution: 1080×1920, screen refresh rate: 60 Hz), positioned 20 cm in front of the mouse. The monitor was slightly angled so that the distance from the mouse to each corner of the screen was equal. Visual trials consisted of directional drifting gratings of 500 ms in duration, and were selected randomly from 6 directions (0, 60, 120, 180, 240, 300 degrees, temporal frequency: 1 Hz, spatial frequency: 0.0450 cycles per degree). Each drifting grating was presented 20 times (total number of trials: 120).

#### Auditory and visual stimuli

Audiovisual trials consisted of pseudo-randomly selected combinations of auditory stimuli and visual stimuli, presented for 500 ms. Each auditory-visual combination was presented 8 times (total number of trials: 1584).

### Electrophysiological recordings

Mice were habituated to head fixation and the experimental setup for approximately 10 days. At least 24 hours before the first electrophysiological recording, animals were anaesthetised using isoflurane (4-5% for induction, ∼2 % maintenance), and a craniotomy of approximately 0.8 mm in diameter was made over A1 (relative to bregma: -2.6 AP, 4.3 ML) using a dental drill, exposing the cortical surface. As before, the skull was covered using two-component silicon putty (Picodent twinsil), over which a hand-cut aluminium cover was glued. Animals were allowed to recover in their home cage for at least 24 hours. On the day of the recording, animals were placed in the experimental setup and head-fixed facing the screen. High-density silicon probes were used to record extracellular activity in A1 (A1x64-Poly2-6mm-23s-160-A64, NeuroNexus, Ann Arbor, USA). The probes had a thickness of 15 µm, and the channels spanned 1449 µm along the length of the probe. A1 was approached from above, using a straight angle relative to the recording table. This resulted in an angle of entry that was not perpendicular to the cortical surface, as well as an approach to A1 at a slight angle (Fig. 1B, C(iii), Fig. S4). We estimated that the resulting angle of insertion was approximately 30° relative to the cortical surface. Due to this angle of insertion, in some sessions, multiple subfields of auditory cortex have been recorded. The probe was slowly inserted (100 µm/min) to a depth of 1500 µm. Once inserted, the probe was allowed to stabilise for at least 10 to 15 minutes before starting the recording. The analogue signal was preamplified and filtered between 0.1 and 20 kHz using an Intan RHD2164 amplifier (Intan Technologies). The Open Ephys Acquisition Board (3^rd^ generation) was used for signal acquisition (continuous sampling rate: 30 kHz) in combination with the Open Ephys GUI^61^

#### Stimulus presentation

During sensory stimulation, all stimulus types (auditory, visual, and audiovisual) were presented to the animal (auditory: 330 trials, visual:120 trials, audiovisual trials: 1584, resulting total number of trials: 2034). Stimulus presentation was pseudo-randomised before the start of each recording and was unique for each session. Inter-stimulus intervals (ISIs) were randomly (normally) distributed between 0.5 and 1.5s in steps of 10 ms. Recordings were performed daily for a maximum of seven days directly following the craniotomy.

### Video monitoring

Orofacial movements were tracked using a monochrome camera (Basler cameras, acA1300-200um, Ahrensburg, Germany) combined with a zoom lens (50 mm, F/2.8 2/3-inch 10 MP, Navitar) positioned approximately 20 cm from the animal, on the left side of the animal. An additional camera was used to track the trunk movements of the mouse (DMK 22BUC03, The Imaging Source, Charlotte, USA), positioned approximately 60 cm from the animal. The face and trunk of the animal were illuminated with an infrared LED (IR-LEDs, 850 nm) to obtain high-contrast illumination of both face and trunk. Animals were placed inside a hollow tube during electrophysiological recordings (Fig. 1A, Fig. S1B); this tube was made of a transparent plastic material to allow for recording of whole-body movement. Orofacial movements were recorded at 100 frames per second (fps) at a resolution of 1280 x 1024. Body movements were recorded at 20 fps at a resolution of 744 x 480. This resulted in a time resolution of 10 ms for the face video and 50 ms for the body video.

### Optotagging: identification of auditory cortex cells projecting to V1

The optotagging protocol was applied to AC cells after all trials with stimulus presentation, so at the end of a recording. The full optotagging protocol was based on single laser pulses of 5 ms, 10 ms, 50 ms, and 100 ms in duration. Pulse durations of these lengths are known to reliably activate ChR2-expressing cells^62,63^. Additionally, to check whether neurons were able to follow high-frequency stimulation, single pulse trials were interleaved with one-second 10 Hz photo stimulus trains of 5 ms laser pulses. The ability to follow high-frequency stimulation supports the classification of a unit as a directly activated and thus, ChR2-expressing cell ^64^. All stimulus types (5 ms, 10 ms, 50 ms, 100 ms, 10 Hz train of 5 ms pulses) were pseudo-randomly presented (each stimulus 25 times), with an inter-stimulus interval (ISI) of 1-1.5 seconds (randomly normally distributed, in 10 ms steps). For light delivery, we used a 473 nm diode-pumped solid-state laser (DPSS, MBL-III-473/1∼100mW, Changchun New Industries, Changchun, China). Light delivery was controlled using a mechanical shutter (LS6, Vincent Associates). A 200 µm core, custom-made multimode optical fibre (FP200URT, Thorlabs) was used for light transmission, coupled to the laser using a 10x objective lens (Thorlabs). The fibre was placed directly over auditory cortex at the start of a recording (Fig. S4).

The full optotagging protocol was performed at two different laser intensities: once with a total power at the tip of 2 mW and once at 5 mW. Only the 2 mW pulses at 10 Hz were used for the classification of tagged units. Four criteria were used to classify neurons as tagged: 1) A significant ZETA test for time-locked modulation in the neuron’s firing rate was required (p-value < 0.01; see *Data analysis,* below); 2) The spike onset latency following laser pulse onset had to be less than 5 ms; 3) Neurons had to fire reliably (i.e., on more than 30% of trials in response to repeated laser pulses, where spikes were counted within a 2–15 ms window after laser onset); (4) The waveform Pearson correlation coefficient between laser-evoked and spontaneous spikes (i.e., preceding laser onset) had to exceed 0.9 (Fig. 1D).

### Histology

Directly following the last recording, animals were euthanised with 20% Euthasol (>1mg/kg; Produlab Pharma, Netherlands) using an intraperitoneal overdose, followed by a transcardial perfusion with 4% paraformaldehyde (PFA, in 0.1M PBS, pH 7.0-7.4). Brains were post-fixed in 4% PFA overnight at 4 °C and then transferred to PBS for temporary storage. Brains were embedded in 3% agarose before sectioning to facilitate cutting of the tissue. Brains were cut in coronal sections using a vibratome (Leica, VT100S) at a thickness of 40 µm and kept in the fridge at 4°C. Sections were then stained with nuclear cell stain (DAPI; 300 nM in PBS, D9542, Sigma) for 1-5 minutes at room temperature (RT), and then washed in PBS three times at RT. Sections were mounted, and an embedding medium (Mowiol; Sigma-Aldrich) was added to preserve fluorescence, together with coverslips. Once the sections were dried, regular nail polish was used to seal the coverslips and the slides, preventing air bubbles. Sections were kept in the fridge at 4°C before imaging. Both viruses used in this study (retrograde-Cre-*tdTomato* and AAV9-*EYFP*) expressed fluorescent reporters; therefore, they did not require secondary antibody staining.

#### Parvalbumin staining

To investigate the possibility of tagged (ChR+) cells also being parvalbumin positive (PV+), for two additional animals, we performed a secondary parvalbumin staining. Tissue was collected, sectioned, and mounted (after the staining) as described above. Sections were washed three times in phosphate buffer (PB, 0.05 M) at RT and then incubated in blocking mix (5 % bovine serum albumin (BSA), in 0.3% phosphate-buffered-saline (PBS)-Triton (PBST)) to prevent nonspecific antibody binding for 2 hours at room temperature. Sections were then incubated with primary antibody (guinea pig anti-PV 1:1000, Synaptic Systems, #195 308) in blocking mix for 1 hour at RT, and then left overnight on a shaker at 4 °C. The next day, sections were washed three times for 10 minutes in 0.3% PBST and then incubated with a secondary antibody (goat anti-Guinea Pig, 1:500, Thermo Fisher Scientific #SA5-10099) and DAPI (H-1200, Vector Laboratories Inc., 1:1000) for 2 hours at RT. Finally, sections were washed for 5 minutes in PBS (0.05 M) and further washed in PB (0.1M) for 5 minutes.

#### Imaging

All sections were imaged at 20x using the slideview VS200 Slidescanner (Evident Scientific) and digitally stored. The second edition of the mouse brain in stereotaxic coordinates^65^ was used as the anatomical atlas in Figure 1 and Figure S5.

### Data analysis

All analyses were performed using Python 3.10; all code used in this article will be made available. The Pynapple package was used for parts of the analysis^66^.

#### Neural data processing

Neural data preprocessing and spike sorting were performed using Spikeinterface^67^. Briefly, the raw neural data were bandpass-filtered between 300 and 6000 Hz. To reduce common noise artefacts, the median of the raw trace of nearby channels (excluding the two directly neighbouring channels in both directions) was subtracted from the bandpass-filtered data. Pre-processed traces were spike-sorted using Mountainsort5^68^. Putative single units were manually curated using Phy GUI (https://github.com/cortex-lab/phy). To further assess the quality of individual units, several quality metrics^69^ were automatically calculated using SpikeInterface^67^. The following cut-off points were used: (1) an inter-spike interval (ISI) violations count < 0.5%; (2) a unit isolation distance^70^ of > 10 and (3) a presence ratio greater than 0.9, indicating that the unit was active across more than 90% of the recording session. We excluded units that had a mean firing rate over the whole session of less than 0.1 Hz.

We also examined whether differences in spike sorting quality metrics between tagged and non-tagged neurons could account for the observed differences in auditory and visual responsiveness. Tagged neurons did not differ in ISI violation count (p = 0.76, two-sided Mann-Whitney U test), but showed small yet statistically significant differences in isolation distance and L-ratio^70^ (isolation distance; p = 0.002, L-ratio, p = 0.01, two-sided Mann-Whitney U test). To assess whether these differences could confound our results, we correlated isolation distance and L-ratio with the outcomes of the auditory and visual ZETA-test p-values of all units (see section below). None of the correlations was meaningful (auditory ZETA vs. L-ratio: r = 0.003; auditory ZETA vs. isolation distance: r = 0.002; visual ZETA vs. L-ratio: r = 0.05; visual ZETA vs. isolation distance: r = -0.08; all p>0.05). This indicates that the slight differences in quality metrics did not contribute to the observed differences between tagged and non-tagged populations in auditory and visual responsiveness.

Firing rates were computed by binning spike times of individual neurons into 10-ms bins. Counts per bin were then smoothed using a causal half-Gaussian window (half-window size of 35 ms). Single-unit firing rates were obtained by dividing the smoothed counts by the length of the window (10 ms). When individual neurons were compared, the firing rate values of each neuron were z-scored per trial. This was achieved by subtracting the mean firing rate of each baseline window (-0.5 to 0 s before stimulus onset) and dividing by the standard deviation of all baseline windows (all trial types).

#### Calculating responsiveness and response latency to laser pulses

To determine whether neurons were responsive to optogenetic stimulation, the recently developed ZETA-test (Zenith of Event-based Time-locked Anomalies) was used^22^ (p < 0.01, Benjamini–Hochberg corrected for false discovery rate ^71^). While the performance of the ZETA test is similar to (or slightly more sensitive than) a more regular t-test, we opted for the ZETA test because it does not require setting arbitrary parameters, such as bin size. We used the minimum duration between individual laser pulses as the analysis window size (0-100 ms after laser onset). Besides determining whether a neuron responds to laser stimulation, the ZETA method is also useful for determining spike response latency to laser onset with high precision. Like the ZETA test itself, it avoids the issue of choosing a bin size, in contrast to peri-stimulus time histogram (PSTH)–based methods. This is achieved by calculating the instantaneous firing rate (IFR) using the temporal deviations upon which the ZETA test is based. Spike response latency is defined as the crossing of the half-maximum response of the peak of the IFR^22^. The peak detection window for determining the latency to optogenetic stimulation was set between 2 and 15 ms.

#### Classification of broad- and narrow-spiking neurons

Neurons were classified into two groups based on two spike waveform characteristics: peak-to-trough delay and peak-to-trough ratio. We fit a Gaussian Mixture Model (GMM) using the ‘Gaussian Mixture’ class from scikit-learn, using these as parameters. The model outputs probabilities of neurons belonging to either of two Gaussian distributions (clusters). Single units that had a probability of less than 85% to belong to either of the clusters were labelled ‘non-classifiable’.

#### Auditory and visual responsiveness

Auditory and visual responsiveness was assessed using the ZETA test over a 0 – 1000 ms window following stimulus onset, with significance defined as p < 0.05. Audiovisual responsiveness required significance in both unimodal conditions. Although the ZETA test is sensitive to both increases and decreases in firing rate, only neurons that increased firing during the analysis window were included in subsequent analyses, ensuring that reported responses reflect stimulus-driven excitation. An exception is the analysis in Fig. 3G-I, where both neurons that either increased or decreased firing rates during visual trials were included, since in this specific case, the interaction between auditory and visual stimulation was studied. Response latency was determined by peak detection within a 5–500 ms window, where the 5 ms lower bound was set based on the minimum response latencies of 5–18 ms reported for primary auditory cortex neurons^23^.

#### Auditory feature tuning

For each neuron, frequency tuning curves were established by calculating the average firing rate during individual trials of a particular frequency and subtracting the mean baseline firing rate, as obtained from all baseline periods, taken as -300 to 0 ms before stimulus onset. The neuron’s ‘best-frequency’ (BF) is defined as the frequency that elicits the maximum response of that neuron during the stimulus period (0 – 300 ms). Only neurons that were significantly responsive to auditory stimuli (ZETA test p < 0.05) were included in this analysis. To compare tuning curves across neurons, each tuning curve was normalised using min-max scaling. For each neuron, the minimum firing rate across all tested frequencies was subtracted from each response, and the result was divided by the difference between the neuron’s maximum and minimum firing rates. This ensured scaling the neurons’ responses in the range of 0-1, allowing comparison across neurons. Furthermore, the responses of each tuning curve were aligned to its best-frequency (BF). The full-width-half-maximum (FWHM) measure was calculated for each neuron by fitting a Gaussian using SciPy curve fit^72^. The fits for each neuron were inspected manually. The same curve-fitting procedure was performed for audiovisual trials to compare auditory tuning in auditory and audiovisual conditions. In a small number of cases (*n =* 5 neurons), the fitting procedure produced non-plausible differences in auditory versus audiovisual conditions; these neurons were therefore excluded from the tuning comparison.

#### Orientation tuning

Significant orientation tuning was estimated using a vector-based approach^27^. For each neuron, mean firing rates were calculated per ‘trial’. A trial is defined here as a single presentation of each stimulus orientation. For this analysis, we used a window between 0 and 300 ms relative to stimulus onset to calculate mean firing rate responses. Since orientation is axis-symmetric, visual stimuli with opposite directions (and thus with the same orientation) were averaged. Next, the responses for individual trials were represented in orientation space (0 to 180°). Orientation vectors were then established per trial by calculating the mean response in orientation space. The Hotelling’s-T² test (a multivariate generalisation of Student’s t-test) was then applied to the distribution of orientation vectors to determine whether they deviated significantly from the origin [0, 0], indicating significant orientation tuning (p < 0.05). The amount of orientation tuning can be quantified using the normalised length of the mean response vector (L_ori,_ or 1-circular variance, see^27^) defined as :

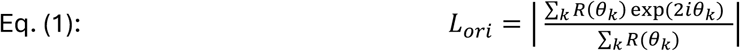

Where *R*(*θ_k_*) is the neural response to the stimuli presented at angle *θ_k_*. This allowed the direct comparison of the tuning of orientation-selective cells in the auditory and audiovisual conditions.

#### Direction tuning

Significant directional tuning was estimated with a vector-based approach^27^. First, the cell’s preferred orientation axis was obtained by calculating the angle of the average orientation vector. For each trial, a direction vector was established by calculating the mean response in direction space. Next, the magnitude of the projection of each direction vector onto the orientation axis was calculated by taking the dot product of each direction vector and the orientation axis (i.e. the ‘direction dot product’). A Student’s t-test was then performed on the distribution of direction dot products. The test determines whether the average magnitude of the distribution of projection vectors is significantly greater than 0, indicating that the cell is significantly direction-selective (p < 0.05).

#### Video analysis

To obtain a quantitative measure of the amount of both orofacial and body movement in the raw videos, FaceMap was used^31^. FaceMap uses singular value decomposition (SVD) on the absolute frame-to-frame differences to quantify the amount of motion (motion energy, ME) in different principal components (PCs) over time. To determine whether an animal was moving or not, we computed the absolute sum of squares (L2 norm) of the first 10 PCs separately for the face (‘face’ video ME) and body videos (‘body’ video ME) (Fig. S1C, D). For plotting and analysis purposes, the video ME of both videos was then binned in 10 ms bins (face video data) or 50 ms bins (body video data) and z-scored by subtracting the mean ME during each baseline window (-0.5 to 0 s before stimulus onset) and dividing by the standard deviation of this baseline window (Fig. S1E, F).

#### Cross-correlation analysis

To assess the possibility of monosynaptic connections between tagged putative excitatory (broad-spiking, BS) and tagged putative inhibitory (narrow-spiking, NS) neurons, a cross-correlation analysis was performed^40^. Cross-correlograms (CCGs) between every neuron pair identified as tagged within a session were computed and compared with a null distribution. This was achieved by computing the number of times a reference cell(e.g. tagged neuron A) fires relative to the other neuron (tagged neuron B) in a particular time-lag bin within a short time window (e.g. -50 to +50 ms). This cross-correlation, the ‘real’ CCG, can be compared with a null distribution by obtaining 1000 ‘jittered’ CCGs by jittering each spike within the spike timestamps of neuron B by a random, normally distributed, amount between 0 and 5 ms. Neuron pairs showing a short, positive latency peak within 1-4 ms, larger than the global 99% band of spike counts, were classified as likely having a monosynaptic excitatory relationship (Fig. S3)^40,41^. Specifically, the global band is constructed by taking the maximum and minimum value of every surrogate (jittered) CCG, and then using the distribution of these extremes to define the upper and lower global thresholds^41^.

#### Mixed-effect model

To account for the hierarchical structure of the data and uneven sampling of neurons across animals, we fit a Bayesian binomial mixed generalized linear model using Statsmodels (BayesianBayesMixedGLM)^73^. Tagged status was used as a fixed effect, with random intercepts and random slopes for tagged status per animal. Parameters were estimated using a variational Bayes mean field approximation (fit_vb function). This analysis was performed separately for auditory, visual and audiovisual stimulus conditions.

#### Logistic regression

To identify single neuron properties associated with auditory and visual responsiveness (ZETA test, p < 0.05), we fit separate logistic regression models for each modality (auditory, visual). Predictor variables included tagged status (tagged, non-tagged), recording depth along the probe (in micrometres), baseline firing rate (Hz), and a binary variable indicating whether the neuron is significantly modulated for the corresponding modality (two-sided Mann-Whitney U test trials with high/low FM, p < 0.05). Models were fit using the Statsmodels Python GLM implementation (model family = *binomial*, link function = *logit*), and coefficients were evaluated using Wald tests to determine statistical significance (p < 0.05). To further investigate the contribution of motor variables to the firing rate of individual neurons in AC, we employed a time-resolved GLM encoding model.

#### Generalized Linear Model

We fit a generalized linear model using the NeMoS package available for Python (https://doi.org/10.5281/zenodo.17553287). The model was trained to estimate the firing rate between -500 and 1000 ms relative to the stimulus onset in 10 ms bins during either auditory or visual trials. We included two sets of predictors to train the model: motor predictors and stimulus-related predictors. For the motor predictors, we used the binned (10 ms bins), z-scored orofacial motion energy (‘face’ video ME), as described previously. The stimulus predictor set consisted of a binary variable indicating whenever a stimulus was present (e.g. “1”, between 0 and 500 ms, and “0” outside this window, -500 to 0 and 500 to 1000 ms relative to stimulus onset), one for each stimulus identity (11 different frequencies for auditory, 6 directions for visual). We presented auditory stimuli at three different decibel levels (65, 70 and 75 dB, SPL), and because firing rates in AC are intensity dependent^23^, we also included stimulus intensity in the (auditory) stimulus predictor set. This was achieved by multiplying the binary stimulus identity by the stimulus intensity on each trial. Furthermore, for both auditory and visual stimulus predictor sets, we used a linearly spaced raised cosine basis function^74^ with 12 basis functions and a window size of 50 bins. These parameters were selected empirically as they yielded a better cross-validated model fit compared to 6 and 10 basis functions. This resulted in a total of 132 (auditory) stimulus predictors and 72 (visual) stimulus predictors. All predictors were z-scored before fitting the model. We fit the model using proximal gradient descent and used elastic net (L1 + L2 regularisation, regularizer strength: 1*10^-4^)^75^.

We fit three models per neuron to estimate the contribution of the different predictor sets to the variance of the neuron’s firing rate: the motor model used only motor predictors, the stimulus model used only stimulus predictors, and the full model used both motor and stimulus predictors. We fit each model for each neuron separately for auditory and visual trials and used five-fold cross-validation. Thus, we split auditory and visual trials into five equal ‘folds’, using four folds for training the model and comparing the model on the held-out fifth part. This procedure was performed five times. Original and predicted firing rates were both smoothed with a Gaussian kernel (standard deviation: 25 ms).

We quantified the contribution of the different predictors to the explained variance (EV) of the neuron’s trial-averaged firing rate using:

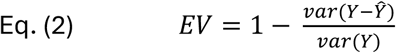

Where *Y* is the original firing rate and *Ŷ* the predicted firing rate. The EV was calculated within the interval 0 – 1000 ms following stimulus onset. This resulted in five EV values per model (one for each fold and per neuron). The EV values presented in Figure 4 represent the average EV over these five folds.

### Statistics

All statistical tests were performed in Python using the SciPy statistical package^72^. To compare two unpaired groups, the two-sided Mann-Whitney U test was used. A one-sided Mann-Whitney U test was used for comparing differences between broad-spiking (BS) and narrow-spiking (NS) single units, as we tested the a priori hypothesis that NS units would show longer spike response latencies, reduced response reliability and higher jitter values due to indirect activation of nearby directly ChR2-expressing neurons (P < 0.05)^39^. The Wilcoxon signed-rank test was used to compare matched pairs of data (e.g., the same neuron across two conditions). As described earlier, for testing individual unit responsiveness to laser stimulation, the ZETA test was used (P < 0.01, Benjamini–Hochberg corrected for false discovery rate ^71^). A more stringent significance threshold was used for laser responsiveness because of its use in unit selection for subsequent analyses. To assess differences in the proportion of auditory- and visual-responsive neurons between tagged and non-tagged groups, neurons were first classified as significantly modulated by auditory or visual stimuli using the ZETA test (P < 0.05). Proportional differences between groups were then evaluated using a two-sided Fisher’s exact test (P < 0.05). The Kolmogorov-Smirnov test (P < 0.05) was used to test whether tagged and non-tagged neurons differed in distributions of best-frequency values.

## Resource availability

### Lead contact

Cyriel M.A. Pennartz

### Materials availability

This study did not generate new unique reagents.

### Data and code availability

Data from this study are available upon reasonable request from the lead contact. Analysis code will be made available on GitLab.

## Acknowledgements

This work was supported by the European Union’s Horizon 2020 Framework Programme for Research and Innovation under the Specific Grant Agreement No. 945539 to C.M.A.P., and by the Netherlands Organisation for Scientific Research (NWO, ENW M2 grant, project OCENW.M20.285) to C.P.J.K and C.M.A.P. We thank Gerjan Huis in ‘t Veld for biotechnical support.

## Author contributions

Conceptualization, S.K. & U.O & C.M.A.P; Data curation, S.K.; Formal analysis, S.K.; Funding acquisition, C.M.A.P; Investigation, S.K.; Methodology, S.K. & U.O & C.M.A.P; Project administration, U.O & C.M.A.P; Resources, U.O & C.M.A.P; Software, S.K.; Supervision, U.O & C.M.A.P; Validation, S.K.; Visualization, S.K.; Writing – original draft, S.K. Writing – review & editing, U.O & C.M.A.P.

## Declaration of interest

The authors declare no competing interests.

## Declaration of generative AI and AI-assisted technologies

During the preparation of this work, the author(s) used ChatGPT version 5.5 in order to improve sentence flow. After using this tool or service, the author(s) reviewed and edited the content as needed and take full responsibility for the content of the publication.

## Supplemental Information

**Document S1. Figures S1-S5.**

## Notes

### Competing Interest Statement

The authors have declared no competing interest.

