## Supplemental Figures S1-S5 for "Visual-cortex-projecting neurons in mouse auditory cortex show distinct audiovisual responses independent of motor-related activity"

### Supplemental Information

**Document S1. Figures S1-S5.**

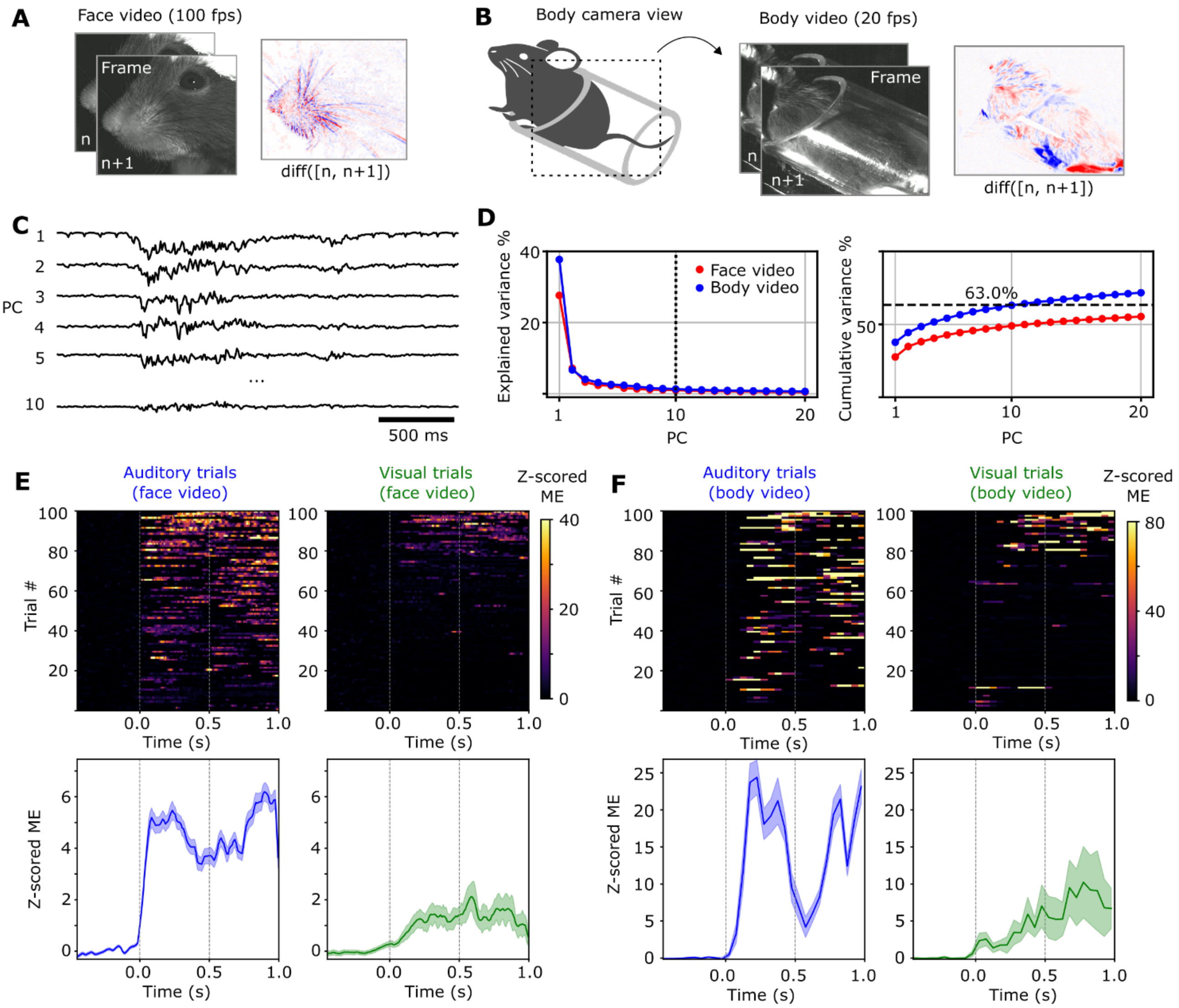


Figure S1. Quantification of orofacial and body movements during auditory and visual trials.

**A**) Orofacial movements were captured at 100 fps, and absolute frame-to-frame differences (diff(n, n+1)) were calculated using FaceMap. **B**) Body movements were captured at 20 fps, and absolute frame-to-frame differences were calculated using FaceMap. The mouse and its enclosure were horizontally positioned. **C**) Absolute frame-to-frame differences were decomposed into 500 principal components (PCs), of which the first 10 PCs are plotted here for an example segment. **D**) Left: the variance explained by each principal component for both the face video (red) and body video (blue). Right: the cumulative variance explained plotted for the first 20 principal components for the face (red) and body video (blue). Cumulative variance explained by the first 10 PCs extracted from the face video data was 48.95% and 63.0% from the body video. These percentages were typical across sessions and animals. **E**) Top panels: face motion energy (face video ME) during auditory (left) and visual trials (right). All trials were individually z-scored and sorted for ME magnitude during the stimulus window (0 – 0.5s); the top 100 trials for each trial type are shown. Bottom panels: average z-scored ME across the trials shown above. Shaded area represents standard error of mean. **F**) Same as plots as in E), now using the z-scored ME from the body video (‘body’ video ME). Trials are sorted using (face) ME magnitude for auditory and visual trials separately. Examples shown here are typical.

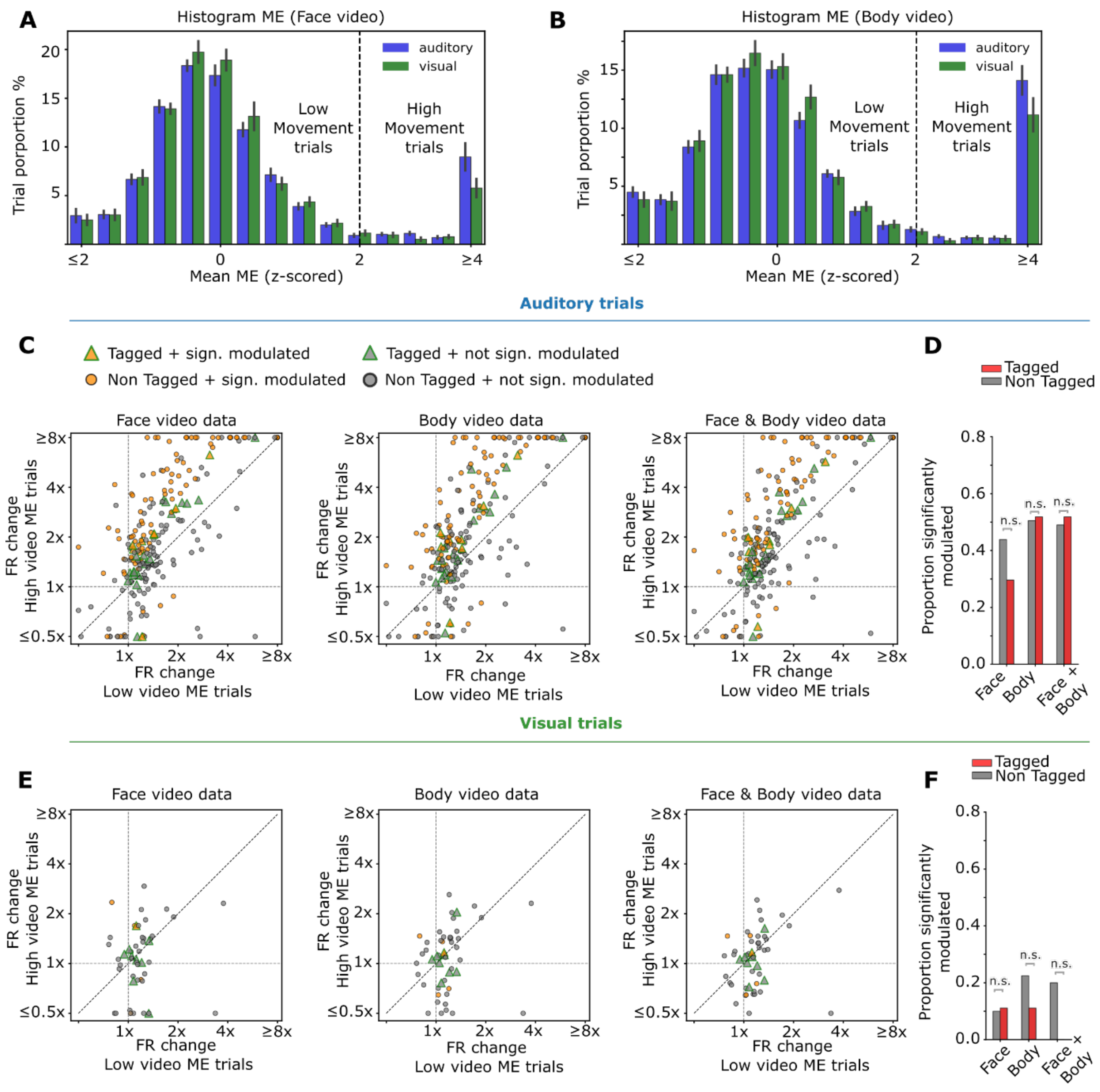


#### Figure S2. Neuronal firing rates were generally higher during auditory trials with high versus low motion energy.

**A**) Histogram of average z-scored ME per trial for auditory and visual trials over all sessions, for all animals (*n* = 13 sessions), calculated using the face video ME. The animal was considered to show a high level of movement whenever the average ME during the stimulus window (0 – 0.5s) crossed 2 standard deviations (SD) above the mean. **B**) Same as A), but we now calculated ME using the body video. **C**) Firing rate (FR) change within the stimulus window (0 – 0.5s) compared to baseline (-0.5 – 0s, relative to stimulus onset) of auditory responsive cells in trials with high orofacial movement (y-axis) and with low orofacial movement (x-axis). To ensure comparability across auditory trials, this analysis was restricted to auditory stimuli with an intensity of 75 dB, as differences in stimulus intensity can also influence firing rate. Middle: same as (left) but now using body ME. Right: same as (left & middle), now classifying trials as 'high' ME whenever either face or body video ME reached > 2 SD. Significantly modulated neurons were classified using face video ME (two-sided Mann-Whitney U test, $p < 0.05$, left plot) and labelled orange in the middle and right plots. Tagged neurons are indicated by a triangular shape. **D**) The proportion of auditory-responsive neurons significantly modulated by movement (tagged vs. non-tagged), separately calculated using the face video ME, body video ME, or face and body video ME combined (Two-sided Fisher's exact test: tagged vs. non-tagged for face, $p = 0.21$, body, $p = 1.00$, face and body, $p = 0.84$, respectively). **E**) Same as plots in C), for visually responsive cells. **F**) Same as D), for visually responsive cells (two-sided Fisher's exact test: tagged vs. non-tagged for face, $p = 1.00$, body, $p = 0.66$, face and body, $p = 0.32$, respectively).

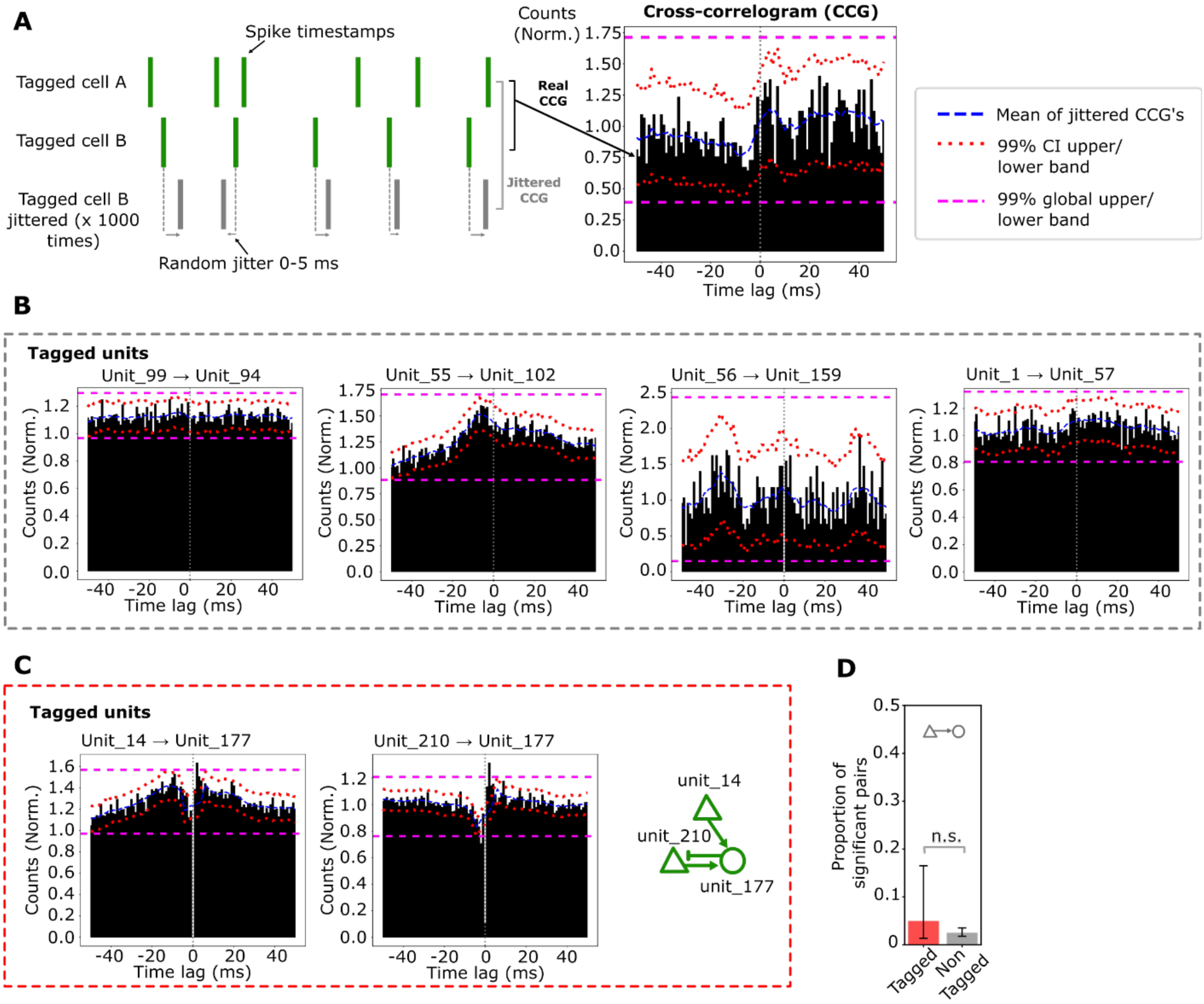


#### Figure S3. Cross-correlation analysis to identify putative monosynaptic connections between tagged neuron pairs.

**A**) Cross-correlograms (CCGs) were computed for all tagged and non-tagged neuron pairs in each recording and compared with 1000 'jittered' CCGs to assess whether tagged neurons were more likely than non-tagged neurons to be monosynaptically connected. Neuron pairs showing a short, positive latency peak within 1-4 ms, larger than the global 99% band of spike counts (purple dashed line), were classified as monosynaptically connected (*See Methods*). **B**) Four example tagged putative excitatory (broad-spiking, BS) to putative inhibitory neuron pairs (narrow-spiking, NS), for which CCGs are shown. In the leftmost example, Unit 99 indicates the reference cell and Unit 94 the target cell. Absence of a short positive latency peak within 1-4 ms does not support monosynaptic connectivity between these tagged neuron pairs. **C**) Two significant tagged neuron pairs (BS→NS) were identified (n = 2 out of 40 total pairs), as indicated by the short positive latency peak crossing the 99% global band of spike counts in both CCGs. These pairs involve the same tagged cell_177.  In green is a schematic drawing of the connectivity between these three cells (Triangles: putative excitatory, Circle: putative inhibitory). **D**) Proportion of all BS→NS cell pairs with a significant positive latency peak within 1-4 ms, larger than the 99% global band of spike counts, for tagged and non-tagged neuron pairs (two-sided Fisher's exact test: tagged vs. non-tagged, p = 0.28).

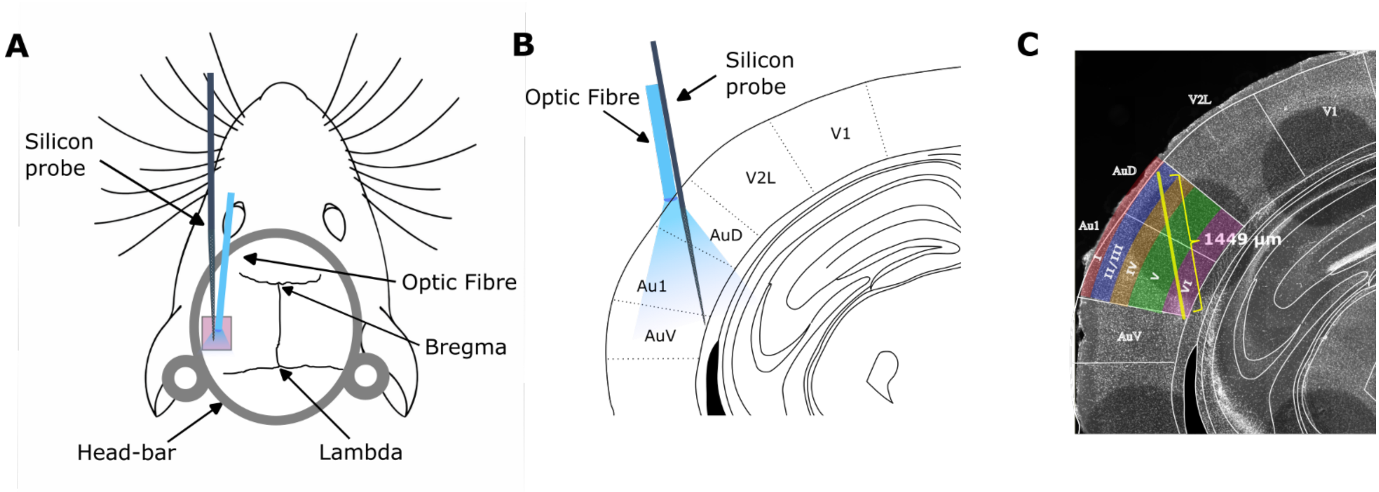


#### Figure S4. Silicon-probe and optic fibre placement during electrophysiological recordings.

**A**) Schematic dorsal view of the mouse in the recording setup. The auditory cortex was approached from above using a straight angle relative to the recording table. **B**) Schematic coronal view of the insertion angle in our recordings. This angle of insertion in B) is obtained due to the slightly lateral placement of the head-bar (see *Methods*) and natural curvature of the brain in this region. **C**) Approximation of the layers of the cortex covered by the full channel span of the silicon probe (yellow line indicates full channel span of 1449 μm). The probe is inserted 1500 μm (see *Methods*). Note that most of the channels (about two-thirds) are situated in layers V and VI, whereas about one-third of the channels cover layers II/III and IV. Exact coverage may slightly vary per recording session/animal. Abbreviations: V1: primary visual cortex, V2L: secondary visual cortex, lateral area. Au1: primary auditory cortex, AuV: secondary auditory cortex, ventral area. AuD: secondary auditory cortex, dorsal area.

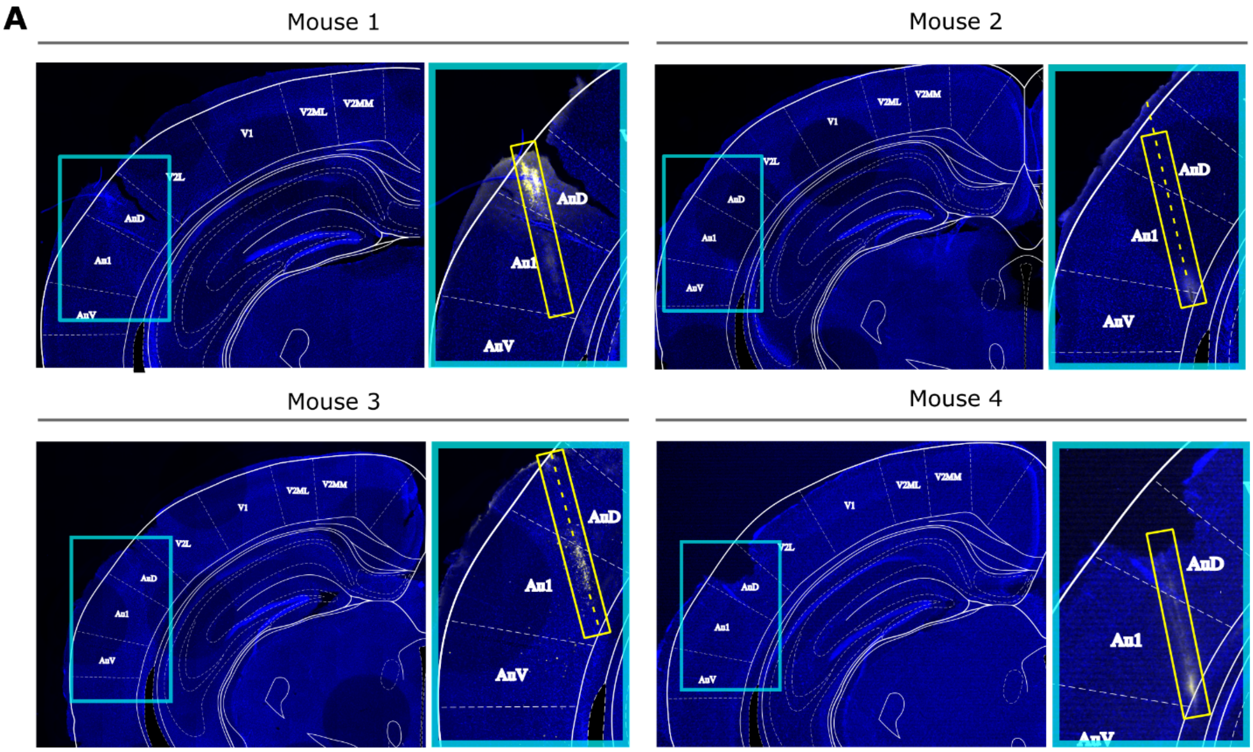


Figure S5. Histological brain sections along with probe tracks for Mouse 1 - 4.

**A**) For each mouse: (left) Hemisphere is manually overlaid with an anatomical atlas (blue: DAPI staining); (right) the electrode track is outlined by the yellow rectangle (yellow staining: CM-Dil dye labelling). In some sections, only part of the probe track is visible, likely due to a sectioning angle that was not parallel to the probe trajectory. In these sections, a dashed yellow line has been manually added to visualise the estimated probe track. The probe tips generally terminated within or just below layer 6. Abbreviations: V1: primary visual cortex, V2MM: secondary visual cortex, medio-medial area. V2ML: secondary visual cortex, mediolateral area. V2L: secondary visual cortex, lateral area. Au1: primary auditory cortex, AuV: secondary auditory cortex, ventral area. AuD: secondary auditory cortex.
